# Regulated compartmentalization of enzymes in Golgi by GRASP55 controls cellular glycosphingolipid profile and function

**DOI:** 10.1101/2020.05.03.074682

**Authors:** Prathyush Pothukuchi, Ilenia Agliarulo, Marinella Pirozzi, Riccardo Rizzo, Domenico Russo, Gabriele Turacchio, Julian Nüchel, Jia-Shu Yang, Charlotte Julie Caroline Gehin, Laura Capolupo, Maria Jose Hernandez-Corbacho, Ansuman Biswas, Giovanna Vanacore, Nina Dathan, Takahiro Nitta, Petra Henklein, Mukund Thattai, Jin-Ichi Inokuchi, Victor W. Hsu, Markus Plomann, Lina M. Obeid, Yusuf A. Hannun, Alberto Luini, Giovanni D’Angelo, Seetharaman Parashuraman

**Affiliations:** Institute of Biochemistry and Cell Biology, National Research Council of Italy, Via P. Castellino, 111, 80131, Italy; Center for Biochemistry, Joseph-Stelzmann-Strasse 52, D50931 Cologne, Germany; Division of Rheumatology, Inflammation and Immunity, Brigham and Women’s Hospital, and Department of Medicine, Harvard Medical School, Boston, MA, USA; École Polytechnique Fédérale de Lausanne (EPFL), AI 1109 (Bâtiment AI), Station 19, CH-1015 Lausanne, Switzerland; Stony Brook University Medical Center, Stony Brook 11794-8430, New York, United States; National Center of Biological Sciences, Bengaluru, India; Division of Glycopathology, Institute of Molecular Biomembrane and Glycobiology,Tohoku Medical and Pharmaceutical University, Sendai, Japan; Universitätsmedizin Berlin Institut für Biochemie Charité CrossOver Charitéplatz 1 / Sitz: Virchowweg 610117 Berlin, Germany

## Abstract

Glycans are important regulators of cell and organismal physiology. This requires that the glycan biosynthesis be controlled to achieve specific cellular glycan profiles. Glycans are assembled in the Golgi apparatus on secretory cargoes that traverse it. The mechanisms by which the Golgi apparatus ensures cell- and cargo-specific glycosylation remain obscure. We investigated how the Golgi apparatus regulates glycosylation by studying biosynthesis of glycosphingolipids, glycosylated lipids with critical roles in signalling and differentiation. We identified the Golgi matrix protein GRASP55 as a controller of sphingolipid glycosylation by regulating the compartmentalized localization of key sphingolipid biosynthetic enzymes in the Golgi. GRASP55 controls the localization of the enzymes by binding to them and regulating their entry into peri-Golgi vesicles. Impairing GRASP55-enzyme interaction decompartmentalizes these enzymes, changes the substrate flux across competing glycosylation pathways that results in alteration of the cellular glycosphingolipid profile. This GRASP55 regulated pathway of enzyme compartmentalization allows cells to make cell density-dependent adaptations in glycosphingolipid biosynthesis to suit cell growth needs. Thus, the Golgi apparatus controls the cellular glycan (glycosphingolipid) profile by governing competition between biosynthetic reactions through regulated changes in enzyme compartmentalization.

## Introduction

Glycans are one of the fundamental building blocks of the cell and play key roles in development and physiology (*1–5*). Cellular glycan profiles are sensitive to changes in cell state and/or differentiation and are also important contributors to the process (*6*). Indeed, several developmental disorders are associated with impaired production of glycans (*7*). Thus, how the glycan biosynthesis is regulated to achieve specific cellular glycan profiles is an important biological problem. In eukaryotes, glycans are assembled mainly by the Golgi apparatus on cargo proteins and lipids that traverse the organelle (*8*). Glycan biosynthesis happens in a template-independent fashion (*9*), yet the products are not random polymers of sugars but a defined distribution of glycans that is cell-type and cargo-specific (*9, 10*). This suggests that their biosynthesis is guided by regulated program(s). Transcriptional programs have been identified that contribute to defining the glycome of a cell, but they only partially account for it (*9, 11, 12*). An obviously important but unexplored factor that influences glycosylation is the Golgi apparatus itself (*13, 14*).

The Golgi apparatus is a central organelle of the secretory pathway that processes newly synthesized cargoes coming from the endoplasmic reticulum (ER), primarily by glycosylation, before sorting them towards their correct destination in the cell. It consists of a stack of 4-11 cisternae (*15*), populated by enzymes and accessory proteins that maintain suitable milieu for the enzymes to act. The stack is polarized with a *cis*-side where cargoes arrive and a *trans*-side from where they leave. The enzymes are not homogenously distributed across the Golgi stack but are restricted or compartmentalized to 1-3 specific cisternae. The cisternal maturation model provides a conceptual framework for understanding Golgi enzyme compartmentalization (*16, 17*). According to the model, secretory cargoes are transported forward by the anterograde flux mediated by cisternal progression, which consists of constant formation and consumption of *cis* and *trans* cisternae, respectively. The retention of Golgi glycosylation enzymes in the face of this continuous forward flux is mediated by their retrograde transport that acts as counterbalance for the forward transport. The retrograde transport is promoted by coat protein complex I (COPI) machinery (*18–22*) and is assisted in this process by adaptor molecules like GOLPH3 (*23–25*), Conserved oligomeric complex (COG) proteins and Golgi matrix proteins especially Golgins (*26–28*). However, the specific molecular mechanisms and processes by which the same retrograde transport pathway promotes localization of enzymes to distinct cisternae remain unknown.

The compartmentalized localization of enzymes has been suggested to influence both sequential as well competing glycosylation reactions. The localization of enzymes along the *cis-trans* axis reflecting their order of action (*29*) has been suggested to influence the efficiency of sequential processing reactions (*30*). On the other hand, the promiscuity of glycosylation enzymes (*31*) makes compartmentalized localization of competing enzymes a critical factor in determining the specificity in glycan output (i.e. the type and quantity of glycans produced) (*29, 32, 33*). When two or more enzymes compete for a substrate, the order in which they get access to it can substantially influence the glycans produced and subsequently the physiological outcomes. Competing reactions are frequent in glycosylation pathways and all known glycosylation pathways have one or more competing glycosylation steps. Nevertheless, how the compartmentalized localization of competing enzymes is achieved, how it is regulated to influence glycosylation reactions and what the physiological relevance of this regulation is remains unexplored.

To evaluate and understand the contribution of Golgi compartmentalization in regulating glycosylation, we have focused our study on sphingolipid (SL) glycosylation. We chose this model system for several reasons: a. it is well characterized from both biochemical and transcriptional perspectives (*6, 34, 35*); b. the glycosylation reaction is less influenced by the cargo structure in contrast to protein glycosylation and thus is a *cleaner* system to study effects of Golgi processes on glycosylation; c. there are simple biochemical methods available to analyse SL glycosylation (*34*) and d. finally SLs have important roles in physiology and development (*36, 37*). The SL glycosylation pathway exhibits the essential features of glycosylation pathways like localization of enzymes reflecting their order of action and also at least two competing reaction steps that are important in determining the metabolic outcome of the pathway (see below). Further, while enzymes of the pathway are well characterized, molecular players regulating their sub-Golgi compartmentalization are unknown. By studying SL glycosylation, we identify GRASP55 as an important factor that compartmentalizes two enzymes catalysing critical branch points of the SL glycosylation pathway. GRASP55 binds to and prevents the entry of these enzymes into retrograde transport carriers. This *retaining* action of GRASP55 is essential for dynamic compartmentalization of these enzymes in the Golgi stack. The competing enzymes thus positioned at appropriate levels in the Golgi stack regulate cargo flux across competing reactions of the pathway and determine the metabolic outcome *viz*. sphingolipid produced by the cell. Further, we find that GRASP55-mediated regulation of SL metabolic outcome controls density-dependent cell growth. These results delineate a molecular mechanism of enzyme compartmentalization and highlight the physiological importance of this mechanism in regulated production of cell surface glycans.

## Results

### Disruption of Golgi organization alters SL biosynthesis

SL biosynthesis starts with the production of ceramide (Cer) in the ER, which is then processed in the Golgi to sphingomyelin (SM) or glycosphingolipids (GSLs). The model cell system we use, HeLa cells, produces two species of GSLs - globosides (Gb3) and gangliosides (GM1 and GM3) (*6, 34, 35*) (See Fig.1A for schematic of the SL system in HeLa cells). This SL pathway includes sequential processing of Cer to complex GSLs as well as two bifurcating steps where the substrates get differentially channelled. The first is the bifurcation between SM and glucosylceramide (GlcCer) biosynthesis, where the substrate Cer is channelled into either of the pathways. The second is the biosynthesis of Gb3 or GM3 from lactosylceramide (LacCer). These two critical steps determine the amount and type of SLs produced by the cell. We first examined the localization of SL biosynthetic enzymes and found that they localize to three distinct zones in the secretory pathway (Fig.1B, **S1)**: (a) the early secretory pathway including the ER and the *cis/medial*-Golgi (C1, C2 cisternae), where Cer biosynthetic enzymes are localized (*33*), have little if any SL biosynthetic enzymes except for a slightly elevated amount of GM3S and GCS in the *cis*/medial-Golgi compared to other GSL biosynthetic enzymes; (b) *medial/trans-*Golgi (C3, C4 cisternae) where most of the GSL biosynthetic enzymes are present alongside substantial amounts of Sphingomyelin synthase 1 (SMS1) and (c) *trans*-Golgi network (TGN), where SMS1 predominates. While all the GSL biosynthetic enzymes show a gradient of increasing concentration from *cis* to *trans*- Golgi, the gradient is much sharper in the case of GB3S and LCS compared to GCS and GM3S **(Fig. S1)**. Thus, the SL biosynthetic enzymes are distributed reflecting their order of action with precursor (Cer) producing enzymes in the early secretory pathway and the Cer processing enzymes in late secretory pathway, which is in turn divided into two distinct zones where GSL and SM biosynthesis predominate. Of note, we expressed HA-tagged enzymes (see Methods) for our studies since the endogenous enzymes were barely detectable and efficient antibodies for EM studies of endogenous enzymes were not available. Nevertheless, the localization mostly reflects expected localization based on enzyme activity and previously published evidence (*38*). A notable exception is the localization of GCS that was shown to be on the *cis*-side of the Golgi (*35*) contrary to what we report here. This is because the earlier studies had used a construct with a tag that blocks the signal for intra-Golgi localization that we identify and describe here. When this signal is blocked, localization of GCS is altered resulting in localization to *cis*-Golgi (see below).

**Figure 1.**
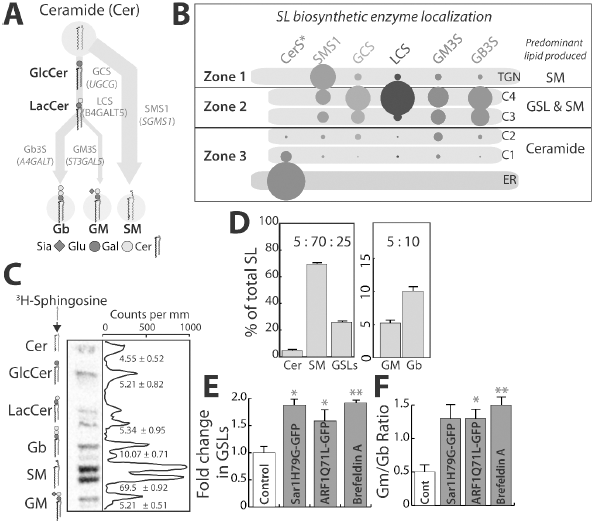
Disruption of SL biosynthetic machinery organization alters SL output: **(A)** Schematic representation of GSL biosynthetic pathway in HeLa cells (Glc, glucose; Gal, galactose; Sia, N-acetylneuraminicacid; Cer, ceramide). Products of biosynthesis are represented in bold and enzymes that catalyse the reactions in grey. The arrows represent the SL metabolic flux from ceramide. **(B)** Schematic representation of GSL biosynthetic zones in HeLa, SM biosynthesis predominates in TGN, whereas GSL and SM production happens in *medial/trans*-Golgi (C3 and C4 cisternae). *Cis*-Golgi/ER is where Ceramide biosynthesis happens with little, if any, SL production. The size of the lipid label arbitrarily represents the proportion of the lipid expected to be synthesized in the compartment based on the localization of corresponding enzymes. **(C)** High-performance thin-layer chromatography (HPTLC) profile of HeLa cells pulsed for 2 hours with [^3^H]-sphingosine and chased for 24 hours. The peaks corresponding to each SL species is indicated and numbers represent each SL species as percentage of total SL. **(D)** The total radioactivity associated with Cer, SM and GSLs (GluCer, LacCer, Gb and GM) or GM and Gb was quantified and presented as percentages relative to total. Data represented as means ± SD of 3 independent experiments. **(E-F)** Biosynthesis of SL in HeLa cells expressing GTP locked mutants of Sar1 or ARF1 or treated with Brefeldin A (BFA; 5μg/ml) was measured by [^3^H] - sphingosine pulse-chase assay. Relative radioactivity associated with Cer, SM and GSLs were quantified and represented as fold change with respect to control. **(E)** For BFA treated cells, the SL output was measured 8h after pulse. **(F)** The ratio of GM/Gb is represented. Data represented as means ± SD of 2 independent experiments. \**P*< 0.05, \*\**P*<0.01 (Student’s t test).

Next, SL output of this system was measured by metabolic labelling with ^3^H-sphingosine, a precursor of ceramide. This revealed the following distribution of products at *quasi steady state* i.e. 24h after labelling: SM (70%), globosides (10%) and gangliosides (5%) and rest remaining as precursors (Cer, GlcCer or LacCer; 15%) (Fig.1C,D). The GSLs (globosides, gangliosides and GSL precursors GlcCer and LacCer) together constituted 25% of total SLs produced. We will refer to the ratio of GSL:SM::25:70 as SL output and the ratio of gangliosides (GM) to Globosides (Gb), GM:Gb::5:10 as GSL output (Fig.1D). For simplicity the SL output will be represented as GSL fraction since a change in GSLs is always associated with a proportional change in SM in the opposite direction. For GSL output, the situation is complex since a substantial portion of signal remains as precursors (GlcCer and LacCer) and so GSL output will be represented as a GM/Gb ratio which under the control conditions corresponds to 0.5 (GM:Gb::5:10). To summarize, SL machinery has a compartmentalized localization across the Golgi in HeLa cells and produces a SL output such that 70% of the Cer is directed towards the production of SM and 25% towards the production of GSLs. Within this 25%, 5% is directed towards the production of gangliosides and 10% towards the production of globosides.

This distribution of glycoforms produced by the Golgi apparatus has largely been ascribed to the expression of the corresponding glycosylation enzymes (*11–13*). To assess the contribution of enzyme compartmentalization to this, we monitored SL output after disrupting the spatial organization of SL biosynthetic enzymes by a) overexpressing GTP locked mutants of monomeric GTPases - secretion associated Ras related GTPase (Sar1 H79G) and ADP ribosylation factor 1 (ARF1 Q71L) that are well-known to disorganize the secretory pathway (*39, 40*) and b) by treating the cells with Brefeldin A, which causes relocation of Golgi enzymes back to the ER. Overexpression of Sar1 H79G led to collapse of the Golgi apparatus into the ER with SL biosynthetic enzymes showing a reticular ER pattern **(Fig. S2A)**. On the other hand, overexpression of ARF1 Q71L mutant led to disruption of stacked cisternal structure of the Golgi, which was replaced by tubulo-vesicular clusters **(Fig. S2B)**, with no separation between *cis* and *trans*-Golgi markers **(Fig. S2C)**. The treatment with Brefeldin A led to the translocation of the enzymes back into the ER as expected, apart from SMS1 which while present in the ER also displayed presence in some punctate structures **(Fig. S2A)**. The SL output was altered in these cells, and consistently in all three conditions there was an increased production of GSLs over SM and gangliosides over globosides **(Fig. S2D,E)**. The SL output represented as fold change in GSL fraction, showed that GSL production in these cells increased by 1.5-1.9 fold over control cells (Fig.1E). Similarly, GSL output measured as GM/Gb ratio changed from 0.5 in control cells to 1.3 - 1.5 in treated cells (Fig.1F). These data suggest that impaired spatial organization of enzymes correlates with altered SL output, and especially the output from steps involving competing reactions are sensitive to disorganization of the Golgi. The contribution of enzyme expression to determination of glycosylation is well established (*11*) but the contribution of the Golgi organization and its importance to this process was not clear. These results underscore a significant and substantive role played by the Golgi apparatus in determining the glycan output of a cell.

### GRASP55 regulates SL output by controlling substrate flux between competing glycosylation pathways

Given the importance of the organization of the Golgi apparatus, and likely of the SL biosynthetic machinery localized to the organelle, to determining SL output, we wanted to identify the molecular players involved in this process. Retention of enzymes in the Golgi depends on their COPI-dependent retrograde transport. Golgi matrix proteins especially Golgins contribute to specificity in this process (*28*) and thus to compartmentalization of enzymes. So, to identify specific regulators of compartmentalization of SL biosynthetic enzymes, we systematically silenced Golgi matrix proteins and studied the effect on SL production. Among the 14 matrix proteins tested by depletion, downregulation of GRASP55 significantly increased the production of GSLs (a 40% increase in GSLs compared to control) while downregulation of GOPC and GCC2 led to a decrease in GSL levels (Fig.2A). We followed up on GRASP55 since its depletion altered SL output similar to that obtained by disorganization of the Golgi apparatus (Fig.1E).

**Figure 2.**
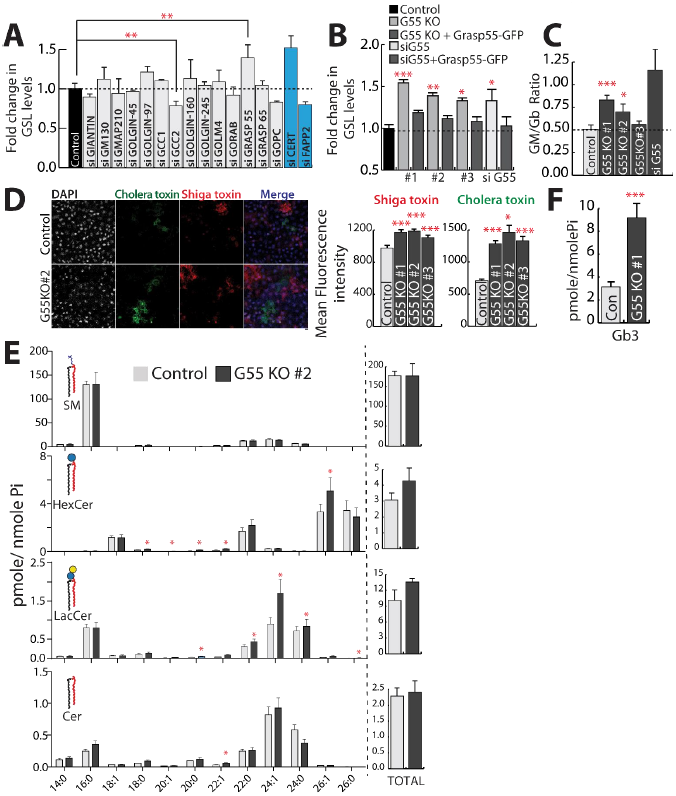
GRASP55 regulates SL biosynthesis: **(A)** HeLa cells were treated with control or indicated siRNA (pool of 4 or 2 as indicated in methods) for 72 hours and SL biosynthesis measured by [^3^H] - sphingosine pulse-chase assay. GSL levels are expressed as fold changes with respect to control. CERT and FAPP2 knockdowns (blue bars) were used as controls. **(B-C)** Effect of GRASP55 depletion on SL biosynthesis monitored by [^3^H] - sphingosine pulse-chase assay in GRASP55 KO cells or cells treated with GRASP55 siRNA or following expression of GRASP55-GFP in GRASP55 depleted cells. GSL levels are expressed as fold changes respect to control. **(C)** The levels of GM and Gb were quantified and represented as GM/Gb ratio. **(D)** Control and GRASP55KO cells were processed for Cy3-conjugated Shiga Toxin (ShTxB) and Alexa488-conjugated Cholera Toxin (ChTxB) staining followed by flow cytometry analysis. Mean fluorescence intensity was measured and represented. **(E-F)** SL levels as assessed by LC/MS or MALDI-MS (Gb3) in control and GRASP55 KO (#2) cells. Data represented are mean ± SD of 3 independent experiments *p <0.05, **p <0.01, ***p <0.001 (Student’s t test).

GRASP55 plays a role in stacking of Golgi *cis*ternae (*41, 42*) formation of Golgi ribbon (*38*), COPI vesicle formation (*43, 44*), secretion (both conventional and unconventional) (*45–47*) and glycosylation (*43, 48*). We generated GRASP55 knockout (KO) HeLa cells (3 independent clones) using clustered regularly interspaced short palindromic repeat (CRISPR)/ CRISPR associated protein 9 (Cas9) technique **(Fig. S3A)** (see Methods). Western blotting and immunofluorescence confirmed the complete abolishment of GRASP55 expression in these KO clones **(Fig. S3B,C)** while expression levels of other Golgi matrix proteins were not altered **(Fig. S3D)**. Fragmentation of Golgi ribbon architecture was confirmed using both *cis* and *trans*- Golgi markers **(Fig. S4A-C)**. The Golgi stack itself did not reveal any obvious alterations **(Fig. S4D)**. Metabolic labelling experiments showed that biosynthesis of GSLs significantly increased in all three clones (by 30-50%) with a corresponding decrease in SM (Fig.2B,**S5A,B**) suggesting a bias towards GSL production in the absence of GRASP55. The total levels of complex GSLs (GM and Gb) also increased and GM/Gb ratio was altered favouring ganglioside production in 2 out of 3 clones (Fig.2C). The increased level of GSLs in cells is also evidenced by increased binding of bacterial toxins (Shiga and cholera toxins that bind Gb3 and GM1 respectively) (Fig.2D,**S5D**). Mass spectrometry analysis showed that there was an increase in the levels of several species of GlcCer, LacCer and globosides in GRASP55 KO cell line (Fig.2E,F). A similar increase in GSLs was also observed in GRASP55 KO human fibroblast cell line (Wi-26) **(Fig. S3E,S5C)**. Finally, the change in SL output associated with GRASP55 abolition was rescued by re-expressing GRASP55-GFP in these cell lines (Fig.2B,**S3C**) suggesting that the observed changes in GSLs were specific. We conclude that GRASP55 acts as a regulator of substrate (Cer and LacCer) flux at the two steps that involve competing reactions – SM *vs* GSLs and gangliosides *vs* globosides in the SL biosynthetic pathway.

### GRASP55 regulates the intra-Golgi localization of GSL biosynthetic enzymes functioning at metabolic branch points

GRASP55 has been proposed to regulate glycosylation by regulating the kinetics of intra-Golgi transport (*43*) and/or ribbon formation (*48*). For GRASP55-mediated regulation of GSL biosynthesis, neither is a likely explanation since the kinetics of GSL biosynthesis **(Fig. S5A)** is very different from that of protein glycosylation (*43*) and downregulation of several matrix proteins known to fragment Golgi ribbon do not affect GSL biosynthesis (Fig.2A). So, to understand how GRASP55 regulates GSL biosynthesis we first studied the consequences of GRASP55 deletion on SM biosynthetic machinery (ceramide transfer protein – CERT and SMS1). GRASP55 deletion did not reduce the levels of CERT or SMS1 **(Fig. S6A,B)**, alter their localization to Golgi **(Fig. S6C,D)** or change the dynamics of CERT **(Fig. S6D,E)**. The kinetics of ceramide transport to the Golgi also remained unaltered **(Fig. S6F)**. These data suggest that SM biosynthesis is not directly affected by GRASP55 deletion.

We next examined the effect of GRASP55 deletion on enzymes of the GSL biosynthetic branch. There were no consistent changes in their levels **(Fig. S7A)** or their presence in the Golgi **(Fig. S7B)** in GRASP55 KO cells. Their intra-Golgi localization was then examined in nocodazole-induced ministacks with GM130 as a marker for *cis*-Golgi/ *cis*-Golgi network (CGN) compartment and TGN46 as a marker for TGN. Nocodazole induced ministacks were used since they show a clearer separation of *cis* and *trans*-Golgi markers and facilitate the intra-Golgi localization of proteins (*49, 50*). The peak localization of GSL biosynthetic enzymes was found in the *medial/trans* part of the Golgi in control cells **(Fig. S8)**. When GRASP55 was deleted the intra-Golgi localization of GlcCer synthase (GCS) and LacCer synthase (LCS) was shifted in the direction of *cis*-Golgi (Fig.3A-B,**S8A-B**) while the localization of SMS1, Gb3 synthase (Gb3S) and GM3 synthase (GM3S) was not altered **(Fig. S8C-E)**.

**Figure 3.**
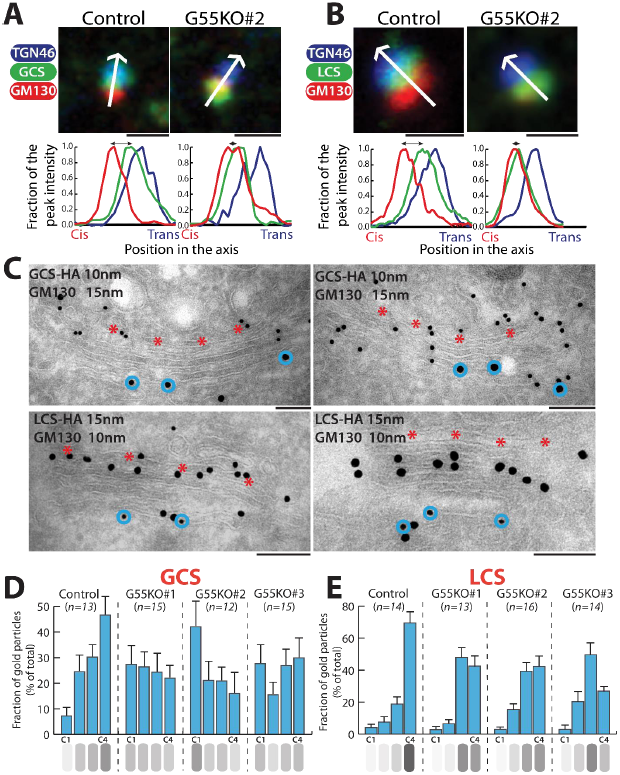
GRASP55 regulates the intra-Golgi localization of GSL biosynthetic enzymes: **(A-B)** Control and GRASP55 KO clones were transfected with HA-tagged GSL biosynthetic enzymes, treated with nocodazole (33 μM) for 3 hours, and processed for immunofluorescence with anti-HA (green), anti-GM130 (red), and anti-TGN46 (blue) antibodies. The relative position of HA-tagged enzymes with respect to GM130 and TGN46 was measured by line scanning and expressed as normalized positions of the peak intensity with the start of GM130 peak indicated as Cis and the end of TGN46 peak indicated as trans in the graph. Scale Bar 1μm. **(C)** Control and GRASP55 KO cells were transfected for 16 hours with GCS-HA or LCS-HA and processed for cryoimmunolabeling with anti-HA antibody (10-nm gold particles) and anti-GM130 antibody (15-nm gold particles) in case of GCS-HA and anti-HA antibody (15-nm gold particles) and anti-GM130 antibody (10-nm gold particles) in case of LCS-HA. Representative images of the distribution of GCS-HA and LCS-HA are shown. Bar 200nm. Red asterisk marks C4 cisterna and blue circles indicate GM130 labelling. **(D-E)** Distribution of indicated enzymes across the Golgi stack was quantified and represented as fraction of Gold particles in each cisternae for GCS-HA **(D)** and LCS- HA **(E)** (n indicated in the graph; data are Mean ± SEM).

The shift towards the *cis*-Golgi was also confirmed by electron microscopy. GCS which is localized mostly to *medial/trans*-Golgi (C3, C4 cisterna) with peak localization in C4 cisterna, became evenly distributed across the stack in GRASP55 KO conditions (Fig.3C,D). The *cis*-most cisterna (C1) which had minimal amount of GCS in control cells showed increased levels of GCS in GRASP55 KO cells, where it reached almost the same level as in other cisternae and in one clone (#2) even resulted in having the peak amount of enzyme (Fig.3D).

In case of LCS, there was a redistribution of the enzyme from *trans*-Golgi (C4 cisterna) to medial-Golgi (C2-C3 cisternae) in GRASP55 KO cells (Fig.3C,E). Unlike GCS, LCS levels in the C1 cisterna did not change significantly (Fig.3E) and thus the shift in intra-Golgi localization was less pronounced in case of LCS. As a control, the localization of SMS1 was not altered under the same conditions **(Fig. S8F)**. To conclude, GRASP55 deletion changed the intra-Golgi localization of two core enzymes of the GSL biosynthetic pathway involved in metabolic branching steps *viz*. GCS and LCS, shifting them from their mainly *trans*-Golgi localization to more *cis/medial*-Golgi localization. These observations raise the following questions: a. What is the mechanism by which the depletion of GRASP55 causes the shift in localization of GCS and LCS? and b. Is the displacement of enzymes responsible for metabolic effects observed after GRASP55 depletion?

### GRASP55 interacts directly with GCS to promote its intra-Golgi localization

To understand how GRASP55 regulates the localization of enzymes we first studied whether GRASP55 interacts with enzymes of the GSL biosynthetic pathway. We expressed GRASP55-GFP and HA-tagged versions of GSL biosynthetic enzymes in HeLa cells, immunoprecipitated GRASP55-GFP and analysed for co-immunoprecipitation of HA-tagged enzymes by western blotting. We found that both GCS and LCS co-immunoprecipitated with GRASP55-GFP while GM3S and Gb3S do not interact. (Fig.4A). The interaction of GCS and LCS with endogenous GRASP55 was also observed in human fibroblast cells (Fig.4B). Surprisingly, in spite of significant homology between GRASP55 and GRASP65 proteins (*41*), the enzymes do not interact with GRASP65, underscoring the specificity of the interaction (Fig.4B).

**Figure 4.**
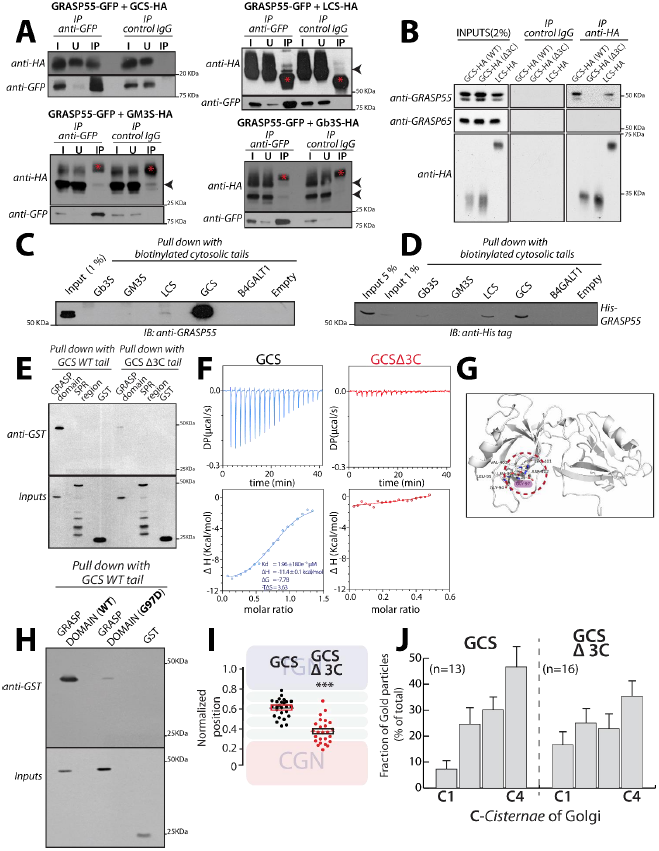
GRASP55 interacts with GCS and LCS: **(A)** HeLa cells co-transfected with indicated HA tagged enzymes (GCS, LCS, GM3S and Gb3S) and GRASP55-GFP were lysed, immunoprecipitated with anti-GFP antibody or control IgG and were analysed by western blotting for interaction by immunoblotting (IB) with anti-HA antibody. I represents 5% on the input lysate, U 5% of the unbound fraction and IP the immunoprecipitate. Red asterisks indicate IgG bands and arrow heads indicates the expected position of HA-tagged enzymes **(B)** WI-26 fibroblasts were transfected with the indicated HA-tagged enzymes were lysed, immunoprecipitated with anti-HA antibody or control IgG and were analysed by western blotting for interaction by immunoblotting with the indicated antibodies. **(C)** Chemically synthesized biotinylated peptides corresponding to cytosolic portions of glycosylation enzymes were bound to avidin beads and were used to pull down interactors from HeLa cell lysates and subjected to immunoblotting with anti-GRASP55 antibody. **(D)** The interaction of chemically synthesized biotinylated peptides, corresponding to cytosolic portions of glycosylation enzymes, with purified His-tagged full-length GRASP55 and their interaction was monitored by pulling down the biotinylated peptides bound to avidin beads followed by western blotting with anti-His tag antibody. **(E)** Chemically synthesized biotinylated peptides corresponding to cytosolic portions of GCS (WT and Δ3C) and indicated purified GST-tagged GRASP domain or SPR region of GRASP55 were incubated together and their interaction was monitored by pulling down the biotinylated peptides with avidin beads followed by western blotting with an anti-GST tag antibody. **(F)** ITC profile, representative of at least 2 independent experiments, for biotinylated GCS and GCS Δ3C cytosolic tails with recombinant GRASP55. **(G)** The molecular basis of interaction between GRASP55 and ceramide glucosyltransferase C-terminal peptide is studied by building a model of GRASP55:GCS peptide structure in the absence of the complex crystal structure. The caroxylate group of Leu of ‘LDV’ motif retains conserved hydrogen bonds with the backbone of 95LLGV98motif of GRASP55. Gly97 residue, which crucial to GRASP:GCS interaction is highlighted (pink). **(H)** The interactions of chemically synthesized biotinylated peptides corresponding to cytosolic portions of GCS (WT) with the indicated purified GST-tagged GRASP domain (WT) or GRASP domain (G97D) were monitored by pulling down the biotinylated peptides with avidin beads followed by western blotting with an anti-GST tag antibody. **(I)** HeLa cells were transfected with either WT GCS or GCS Δ3C, treated with nocodazole (33 μM) for 3 hours and labelled for enzymes, GM130, and TGN46. Line scan analysis was performed as in Fig. 3A-B and the relative position of enzymes was quantitated and plotted. The data are mean ± SD; n> 30 stacks. ***p <0.001(Student’s t test). **(J)** HeLa cells were transfected with either WT GCS or GCS Δ3C for 16 hours with GCS-HA or LCS-HA and processed for cryoimmunolabeling. Distribution of indicated enzymes across the Golgi stack was quantified and represented as fraction of Gold particles in each cisterna. (n indicated in the graph; data are Mean ± SEM).

GRASP55 is a peripheral membrane protein that is anchored to the Golgi membrane through its myristoylated N-terminus. So, interaction with GRASP55 is likely mediated by cytosolically exposed portions of the enzymes. Many Golgi glycosylation enzymes are type II membrane proteins with a short N-terminal cytosolic tail, a transmembrane domain and the luminal enzymatic domain. Thus, the likely part to interact with GRASP55 is the short N-terminal tail. GCS is an exception to this rule and is a multi-transmembrane protein with both N and C-terminal cytosolic tails and a large catalytic portion facing the cytosol. We noticed that the C-terminal tail of GCS ends with a consensus class II PDZ domain interacting motif (Φ-X-Φ-COOH, where Φ refers to hydrophobic amino acid and X to any amino acid) that can potentially bind to the N-terminal tandem PDZ domains of GRASP55. We chemically synthesized the C-terminal cytosolic tail of GCS and N-terminal cytosolic tails of other GSL biosynthetic enzymes and studied their interaction with GRASP55 present in cell lysates as well as purified His-tagged protein.

We found that GCS tail strongly bound to GRASP55 from both cell lysates and the purified protein unlike the tails of other GSL biosynthetic enzymes (Gb3S or GM3S) or the tail of B4GALT1 (see below for description of LCS binding) (Fig.4C,D). We used B4GALT1 a galactosyltransferase involved in protein glycosylation since it was unrelated to the GSL biosynthetic pathway. GRASP55 consists of two domains – N-terminal tandem PDZ domains (GRASP domain) followed by a serine proline rich (SPR) region (*30*). The cytosolic tail of GCS interacted with GST-tagged GRASP domain of GRASP55 but not with GST-tagged SPR region (Fig.4E). Deleting the Φ-X-Φ-COOH motif in GCS cytosolic tail (GCS-Δ3C tail) impaired its interaction with GST-tagged GRASP domain of GRASP55 (Fig.4E). Further, deleting this motif also impaired the interaction of full-length GCS (GCS-Δ3C) with GRASP55 in cells as evidenced by co-immunoprecipitation analysis (Fig.4B). Next to gain further insights into this interaction, we studied it by using isothermal titration calorimetry (ITC). By ITC, we found that cytosolic tail of GCS interacts strongly with the GRASP domain of GRASP55 with a K_d_ value of 2μM (Fig.4F). Of note, the K_d_ values of interaction between GRASP55 and the Golgin-45 tail was approx. 0.27μM (*51*) and that of the LCS tail and GOLPH3 (another Golgi matrix protein that interacts with Golgi enzymes) is approx. 60μM (*23*). There was no significant interaction of GRASP domain of GRASP55 with GCS tail deleted of Φ-X-Φ-COOH motif or B4GALT1 tail (Fig.4F). These studies in sum suggest that GCS C-terminal tail directly and specifically interacts with the GRASP domain of GRASP55.

To understand the amino acid residues in GRASP55 that are critical for this interaction we resorted to modelling. The crystal structure of the GRASP domain of GRASP55 bound to Golgin45 C-terminal region (which contains a hydrophobic amino acid similar to GCS) showed that the peptide binds to a cleft between the PDZ1 and PDZ2 domains of the proteins (*51*). A conserved pocket in the GRASP domain consisting of ^95^LLGV^98^ corresponding to X-Φ1-G-Φ2 motif (X: any amino acid, Φ: hydrophobic amino acid, G: glycine), acts as a binding site for the C-terminal end of the interacting peptide forming hydrogen bonds with the last four residues of Golgin-45 peptide. The X-Φ1-G-Φ2 peptide has a strained left-handed helix conformation, which is usually populated by glycine residues in the Ramachandran plot (*52*). B-factor analysis of the crystal structure of the GRASP55:Golgin-45 complex, revealed that the X-Φ1-G-Φ2 motif forms a rigid loop **(Fig. S9A)** where usually Gly is favoured over Asp. Indeed, substitution of Gly to Asp abolished the GRASP55-Golgin-45 peptide interaction (*51*). A model of GCS peptide was first built using backbone conformation of Golgin4-5 peptide as a template and docked onto the cleft between PDZ1 and PDZ2 domains. The lowest energy model generated was analysed to probe the protein:peptide interaction. The structural analysis also indicated the important contribution of the ^95^LLGV^98^ pocket to binding with the C-terminal amino acids of the GCS peptide (Fig.4G). The predicted binding affinity (ΔG) between protein and peptide was –13.1 kcal mol^-1^, indicating a favourable interaction and when ‘LDV’ was removed from the peptide, the binding affinity was reduced (ΔΔG = −1.5 kcal mol^−1^) indicating the importance of the backbone mediated conserved hydrogen bonds in peptide binding. Given the overall similarity between the GRASP55-Golgin-45 peptide and GRASP55-GCS peptide interactions, we tested if the interactions show similar sensitivities to mutations. As mentioned before the GRASP55-Golgin-45 peptide interaction is sensitive to the substitution of Gly^97^ to Asp (*51*) and so we introduced the corresponding G97D mutation in the GRASP55 PDZ domain to examine its effect on the interaction with the GCS tail. The interaction of the mutant protein GCS C-terminal tail was greatly diminished (Fig.4H). Thus, the interaction between GCS and GRASP55 is likely mediated by the C-terminal LDV motif of GCS interacting with the ^95^LLGV^98^ pocket in the GRASP domain of GRASP55 with Gly97 playing a critical role in the process (Fig.4H). Next, we examined if binding to GRASP55 is essential for intra-Golgi localization of GCS. The last 3 C-terminal amino acids of GCS were deleted (GCS-Δ3C) and the localization of the mutant protein was studied. The intra-Golgi localization of GCS-Δ3C was altered, with it displaying a more *cis*-Golgi localization compared to the WT enzyme (Fig.4I,J). Thus, direct interaction with GRASP55 is essential for the correct sub-compartmentalization of GCS.

Compared to GCS, the LCS tail showed a qualitatively weaker interaction with GRASP55 both for endogenous and recombinant protein (Fig.4C). Nevertheless, it was significantly above that of other GSL biosynthetic enzymes (GM3S or GB3S) and that of B4GALT1. This weak interaction was mediated by the GRASP domain of GRASP55 **(Fig. S9B**). Nevertheless, ITC studies did not show any significant interaction between purified LCS cytosolic tail and the GRASP domain of LCS **(Fig. S9C)**. Thus, in the case of LCS, while the full-length enzyme co-immunoprecipitated with GRASP55, the cytosolic tail of the enzyme itself shows only a weak or no interaction with the protein suggesting that the interaction between LCS and GRASP55 is likely to be indirect. The change in localization of LCS in the absence of GRASP55 correlates with observed interaction with GRASP55 (Fig.4A,B).

In summary, we conclude, that GCS and LCS interact with GRASP55 and in the case of GCS this interaction is direct and is essential for its intra-Golgi localization.

### GRASP55 compartmentalizes the enzymes by preventing their entry into retrograde carriers

According to the cisternal maturation model, the retention of resident proteins in the Golgi is due to their continuous retrograde transport mediated by COPI in the face of anterograde flux of cargoes (*16, 17*). In the framework of this model, compartmentalization of enzymes to specific cisternae of Golgi is achieved by a balance between the anterograde and retrograde flux of enzymes. Indeed, impairing the retrograde transport of enzymes promotes their forward transport leading to their localization to post-Golgi compartments (*50*). So, to understand how this balance is affected in GRASP55-depleted cells to change the steady state localization of GCS and LCS, we examined the distribution of GCS and LCS in peri-Golgi vesicles/carriers. These vesicles depend on COPI for their formation and entry into these vesicles is essential for the intra-Golgi retrograde transport of proteins (*50*). In control HeLa cells, the distribution of enzymes in peri-Golgi vesicles varied. For instance, the density of LCS in peri-Golgi carriers was nearly the same as that of the cisterna while the density of GCS in vesicles was 1.8-fold more than that of the cisterna. Further SMS1, an enzyme unaffected by GRASP55 depletion, was depleted in peri-Golgi vesicles compared to cisterna (density in vesicles was 0.3-fold that of the cisterna) (Fig.5A-C). These differences in density of enzymes in vesicles correlate well with their observed distribution in the Golgi at steady state *i.e.* an increased presence in peri-Golgi vesicles correlates with increased *cis*/medial-Golgi localization. For instance, GCS with higher relative density in peri-Golgi vesicles also has a higher *cis*-Golgi to TGN ratio than SMS1, which has a lower relative density in peri-Golgi vesicles **(FIG. S1)**. This suggests that the presence of proteins in retrograde transport carriers could be a reliable indicator of intra-Golgi distribution of proteins. Next, we studied the density of enzymes in peri-Golgi vesicles in the absence of GRASP55. We found that the density of GCS in vesicles increased to 2.5-fold compared to that present in the cisterna and that of LCS increased to 2-fold over that of the cisterna, while that of SMS1 was unaltered (Fig.5A-C). Thus, GCS and LCS that show an increased localization to cis/medial-Golgi in the absence of GRASP55 also show an increased presence in vesicles in GRASP55 KO cells compared to control, while vesicle distribution of SMS1, whose intra-Golgi localization is unaltered by the absence of GRASP55, remains unchanged. To further validate the increased entry of GCS and LCS into peri-Golgi carriers in the absence of GRASP55, we resorted to *in vitro* budding assays to purify COPI coated vesicles from Golgi membranes using well-established methods (*53*). The Golgi apparatus was purified from control and GRASP55 KO (#2) cells and were incubated with purified coatomer, myristoylated ARF1 and BARS (Brefeldin A ADP ribosylation substrate) to promote budding of COPI retrograde carriers from the Golgi. The budded vesicles were then separated from the Golgi apparatus by centrifugation. The Golgi apparatus was recovered in the pellet fraction while purified COPI vesicles remained in the supernatant fraction **(Fig. S10A)**. We then analysed the presence of LCS and GCS in the supernatant fractions by western blotting. We found that there were increased amounts of LCS and GCS in COPI vesicles that budded from the Golgi apparatus purified from GRASP55 KO cells as compared to control cells (Fig.5D,E), thus confirming our observations with EM. This specific increase in amounts of GCS and LCS in the peri-Golgi carriers in the absence of GRASP55 is consistent with the hypothesis that GRASP55 limits the retrograde transport of these enzymes.

**Figure 5.**
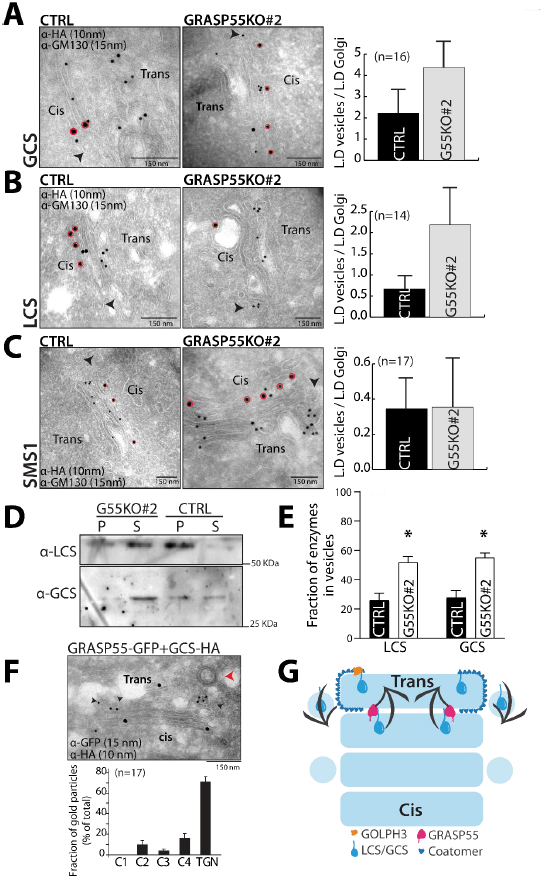
GRASP55 compartmentalizes the enzymes by preventing their entry into retrograde carriers: **(A-C)** Control and GRASP55 KO (#2) cells were transfected for 16 hours with the indicated HA-tagged enzymes and processed for cryoimmunolabeling with anti-HA antibody (10-nm gold particles) and anti-GM130 antibody (15-nm gold particles). Representative images of the distribution of HA-tagged enzymes are shown. Red circles indicate GM130 labelling. Arrow heads represent the peri-Golgi vesicles. Bar 150 nm. Quantification of the distribution of enzymes in vesicles represented as normalized linear density (n indicated in the graph; data are Mean ± SEM). **(D)** COPI vesicles were reconstituted using the two-stage incubation system (detailed in the Methods section). After the second-stage incubation, samples were centrifuged to obtain the pellet fraction that contains Golgi membranes and the supernatant fraction that contains reconstituted COPI vesicles. Both fractions were immunoblotted for LCS and GCS to show their relative distributions on Golgi membranes and in COPI vesicles. **(E)** The COPI vesicle reconstitution system was performed as described above (in D), and then the fraction of LCS and GCS in COPI vesicles versus their total distribution (on Golgi membranes and in COPI vesicles) was calculated. The mean and standard error from 3 independent experiments are shown, * p < 0.05, Student’s t-test. **(F)** HeLa cells co-transfected with HA-tagged GCS and GRASP55-GFP were incubated for 16 hours and were processed for cryoimmunolabeling with anti-HA antibody (10-nm gold particles) and anti-GFP antibody (15-nm gold particles). Representative images of the distribution of GCS and GRASP55-GFP are shown. Red arrowhead indicates the clathrin-coated vesicle that marks the TGN area and black arrowheads indicate the presence of GCS-HA in TGN. Bar 150 nm. Distribution of GCS across the Golgi stack and TGN was quantified and represented as fraction of gold particles. (n indicated in the graph; data are Mean ± SEM). **(G)** Model represents GRASP55-mediated compartmentalization of GCS and LCS. A cyclical and balanced activity of GRASP55 and GOLPH3 compartmentalizes LCS/GCS to the trans-Golgi. The anterograde transport of enzymes (forward direction arrow; cis to trans direction) counterbalances their retrograde transport (reverse direction arrow; trans to cis direction) resulting in the compartmentalization of these enzymes.

If absence of GRASP55 alters the distribution of the protein, does an increase in GRASP55 levels also influence their distribution? To test this, we overexpressed GRASP55-GFP in cells along with GSL biosynthetic enzymes LCS and GCS. In the case of LCS, when GRASP55 was overexpressed, there was a change in localization of the enzyme, which was now present in endosome-like structures along with the Golgi **(Fig. S10C)**. These structures were similar to those to which LCS localizes in the absence of GOLPH3, an adaptor that links LCS to COPI and thus promoting its retrograde transport through peri-Golgi vesicles (*46*). This suggests, that overexpression of GRASP55 likely inhibits retrograde transport in a way that is similar to the absence of GOLPH3. On the other hand, overexpression of GRASP55 did not shift GCS to a post-Golgi localization **(Fig. S10B)** but GCS was increasingly found in the TGN under these conditions (Fig.5F). This suggests that GCS behaves in a similar way to LCS when GRASP55 is overexpressed *i.e.* its localization shifts to a forward position along the secretory pathway. This distribution of enzymes to TGN or post-Golgi compartment when GRASP55 is overexpressed is consistent with an impairment of their retrograde transport. Thus, GRASP55 likely acts to inhibit retrograde transport of LCS and GCS, such that in its absence the enzymes shift to a *cis*/medial-Golgi localization and when GRASP55 levels are increased they shift to a TGN/post-Golgi localization.

We propose that GRASP55 inhibits retrograde transport of LCS and GCS by acting as a “retainer” that binds to and prevents their entry into retrograde transport carriers. While the action of adaptors, exemplified by GOLPH3, that promote entry into retrograde transport vesicles is well accepted, a *retainer* action in the Golgi apparatus has not been hypothesized before. A retainer molecule that prevents the entry of its interactors into peri-Golgi vesicles can be expected to be absent from the peri-Golgi vesicles unlike adaptor molecules (GOLPH3) that promote the sorting of their interactors into peri-Golgi vesicles (*27*). So, we analysed the distribution of GRASP55 on the cisterna and on peri-Golgi vesicles. We find that while nearly half of the Golgi-localized GOLPH3 was found in peri-Golgi carriers only about 8% of the Golgi-associated staining of GRASP55 is associated with peri-Golgi vesicles, suggesting that GRASP55 is likely excluded from peri-Golgi vesicles **(Fig. S10D)**.

Rationalizing, these observations into a model we propose that recycling adaptors and retainers act in an opposing manner to promote compartmentalization of enzymes in the Golgi apparatus. The binding to COPI either directly (*19*) or through recycling adaptors like GOLPH3 (*24, 25*) promotes the retrograde transport of enzymes thus preventing their exit from the Golgi. On the other hand, retainers bind to their client molecules to prevent their entry into retrograde transport carriers and thus indirectly promote their anterograde transport by cisternal progression (Fig.5G). Thus, a cyclical and balanced transport of enzymes – in the retrograde direction by COPI machinery assisted by recycling adaptors and in the anterograde direction by cisternal flow assisted by retainers, compartmentalizes them to specific cisterna of the Golgi apparatus.

### Change in intra-Golgi localization of GSL biosynthetic enzymes changes GSL output

We then examined if the change in localization of enzymes following reduction in GRASP55 levels contributes to the associated changes in GSL biosynthesis. When GSL biosynthesis was examined in Brefeldin A treated cells, the increased GSL production in GRASP55 KO cells as well as the increased GM/Gb ratio were not observed (Fig.6A,B), suggesting that compartmentalized localization of enzymes is essential to manifest GRASP55-deletion induced alteration in GSL production. Next, we examined GSL biosynthesis after expression of GCS and LCS mutants that have altered intra-Golgi localization. We expressed three GCS mutants: a. GCS-Δ3C that shows more *cis*-Golgi presence than the WT enzyme (Fig.4J), b. GCS with HA-tag at C-terminus (GCS-HA_C_) that is expected to block accessibility to the C-terminal valine. GC-SHA_C_ distributes across the stack with significant presence in *cis*-Golgi (*30*) and unlike GCS-Δ3C it was also partially localized to ER **(Fig. S11A)** and c. GCS with HA-tag in the N-terminal tail (GCS-HA_N_), which localizes to ER **(Fig. S11A)**. Thus, they have distinct but overlapping distribution along the secretory pathway. Expression of the wild type GCS construct in HeLa cells led to a 10% increase in the total GSLs produced compared to non-transfected cells suggesting that expression of the construct does not overwhelm the biosynthetic system. The expression of GCS-Δ3C, which localizes to the *cis*-Golgi unlike wild type GCS, led to a 30% increase in GSLs produced, a difference that is quantitatively similar to what is observed between control and GRASP55 KO cells. This suggests that the increased production of GSLs under GRASP55 KO conditions can be explained by the shift in localization of GCS. Interestingly, an increasing ER localization of GCS (GCS-HA_C_ and GCS-HA_N_) led to a larger increase in GSL production (2.3 and 2.6-fold increase respectively) (Fig.6C). The observed differences in GSL production are not due to differences in expression since addition of Brefeldin A neutralized these differences (Fig.6C), implying that changes in localization were indeed the cause for the observed increase in GSL production.

**Figure 6.**
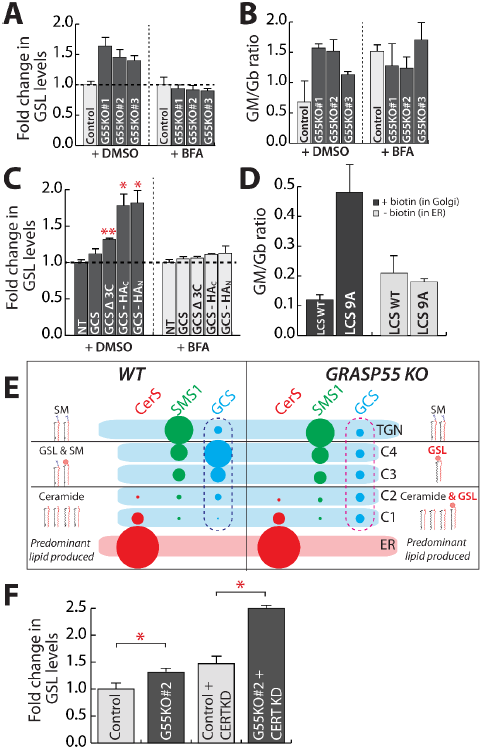
Change in intra-Golgi localization of GSL biosynthetic enzymes changes GSL output: **(A-B)** Control and GRASP55 KO clones were pre-treated with DMSO or BFA (5μg/ml) for 30 min and SL output monitored by [^3^H] - sphingosine pulse-chase assay (8 h chase). Total GSL levels were quantified and expressed as fold changes with respect to control **(A)**. The GM/Gb ratio was calculated and represented **(B)**. Data are mean ± SD of 2 independent experiments. **(C)** HeLa cells were transfected with indicated GCS constructs for 16 hours and SL output monitored by [^3^H] - sphingosine pulse-chase assay (8 h chase) in the presence or absence of BFA (5μg/ml). Total GSL levels were quantified and expressed as fold changes respect to control. Data are mean ± SD of 2 independent experiments. **(D)** HeLa cells KO for LCS were transfected with indicated LCS RUSH constructs for 16 hours, and expressed enzymes were retained in the ER (-biotin) or placed in the Golgi by addition of biotin (40 μM) (+biotin) and SL output monitored by [^3^H] - sphingosine pulse-chase assay. The ratio of GM/Gb is represented. Data are mean ± SD of 2 independent experiments. **(E)** Schematic representation of how the change in localization of GCS in GRASP55 KO with respect to control (pink dashed box vs blue dashed box) from trans- to cis-Golgi results in preferential access to ceramide and thus an increased production of GSLs. (F) CERT was silenced using siRNAs for 72 hours in control and GRASP55 KO (#2) clone before subjecting to [^3^H] –sphingosine pulse chase assay. GSLs were quantified and represented as fold change with respect to control. Values are mean ± SD (n=3).

Next, we examined if the next branch point in the SL metabolic pathway *viz* GB3-GM3 branch was also sensitive to enzyme localization. The key enzyme regulating this branch point is LCS, whose localization is again controlled by GRASP55. While we find that GRASP55 and LCS interact, there is no convincing evidence for their direct interaction so mutating key residues to decompartmentalize the protein similar to what was achieved for GCS is not possible. While studying the molecular basis of LCS localization it was found that the 14 amino acid cytosolic tail of LCS contains the information needed to localize the protein in the Golgi (*23*). So, we performed alanine mutagenesis of most of the cytosolic tail of LCS (LCS9A) except for the membrane proximal 5 amino acid region essential for interaction with GOLPH3 and the subsequent retention of LCS in Golgi. We found that while LCS9A was still retained in the Golgi apparatus as expected, it localized in a de-compartmentalized manner with no clear trans-Golgi localization as observed with LCS WT **(Fig. S11B).** We do not know the reason why these mutations lead to a de-compartmentalized localization of LCS, but they provide an opportunity to test whether compartmentalization of LCS regulates flux across Gb3-GM3 branches similar to what was observed with GCS. To this end, we analysed the GSL output following the expression of LCS WT and LCS9A constructs in LCS KO HeLa cells (*54*). We found that the expression of LCS9A in Golgi favoured the production of gangliosides over globosides compared to LCS-WT. Thus GM/Gb ratio of 0.12 that is observed in case of LCS-WT expressing cells changed to 0.47 in case of LCS9A expressing cells (Fig.6D). The differences were nullified when the enzymes were retained in the ER (Fig.6D) suggesting that the observed differences were likely due to altered intra-Golgi localization of these enzymes. Thus, a change in LCS localization observed in GRASP55 KO conditions may contribute to the observed increase in GM/Gb ratio in these cells. From these data we conclude that altered localization of key enzymes of the GSL pathway can reproduce the effects on SL biosynthesis following GRASP55 deletion and that localization of enzymes in the Golgi can control cargo flux across competing biosynthetic pathways.

To explain the effects of GRASP55 KO on GSL biosynthesis based on these results, we thus considered the localization of these respective enzymes. As discussed earlier, most of the GSL biosynthesis likely happens in the medial/*trans*-Golgi where along with the GSL enzymes a substantial portion of SMS1 is also located (Fig.1B). This likely leads to a competition between SMS1 and GCS for Cer that is transported by vesicular transport to the *medial/trans*-Golgi (Fig.6E). By moving GCS to the *cis*-Golgi, as happening in GRASP55 KO cells, the enzyme now gets a preferential access to Cer and resulting in an increased production of GSLs (Fig.6E). Indeed, in CERT KD cells where the non-vesicular transport of Cer is blocked and Cer reaches the Golgi mostly by vesicular transport, the effect on GSL biosynthesis resulting from the absence of GRASP55 is further increased (Fig.6F) suggesting that GCS and SMS1 compete for Cer that is transported by a vesicular transport pathway to the Golgi. The effect of GRASP55 KO on GSL output can also be explained by a similar logic. While both GM3S and Gb3S are localized mainly to the *medial/trans*-Golgi, significant amounts of GM3S can also be seen in *cis/medial*-Golgi unlike Gb3S **(Fig. S1)**. Thus, when LCS is moved towards the *cis/medial*-Golgi, ganglioside biosynthesis by GM3S is favoured over globoside biosynthesis likely due to preferential access of GM3S to the substrate LacCer.

Thus, these data provide a model of how compartmentalization of enzymes in the Golgi contributes to faithful glycan output. When two enzymes compete for a common substrate and are localized in the same compartment, the product of a glycoenzyme with more affinity for substrate and/or increased expression dominates the glycan output. On the other hand, if one of the competing enzymes is present in an earlier part of the secretory pathway relative to the other, it gets preferential access to the substrate before its competitor and thus promotes flux across the pathway catalysed by it resulting in a glycan output that has a bias towards the product of this enzyme. Thus, positioning of enzymes in the Golgi regulates the pattern of glycan distribution in the output.

### GRASP55 contributes to confluency induced changes in GSL production

We then explored the physiological relevance of this GRASP55-mediated regulation of GSL biosynthesis. Confluence of fibroblasts is known to regulate GSL biosynthesis, which is required for cell density induced contact inhibition of growth (*55*); so we tested if this involves GRASP55-mediated regulation of GSL biosynthesis. We cultured human fibroblasts (wi-26) to different extents of confluence (25% to 100%) (Fig.7A) and monitored SL biosynthesis. A change in cell confluence was accompanied by a change in SL output. Going from sparse (25%) to complete confluence (100%), there was an increased production of SM with a corresponding decrease in GSL biosynthesis. SL output measured as GSL levels showed that there was an approximately 20% reduction in GSL production in completely confluent fibroblasts compared to sparse conditions (Fig.7B). Next, we examined the reason behind the change in production of GSLs when cells reach confluence. It has been suggested that expression of GSL enzymes is altered with confluence (*55*). So, we analysed the levels of the SL biosynthetic enzymes in sparse and confluent cells by quantitative RT-PCR. As published earlier, we also found that the enzymes belonging to the distal end of GSL biosynthetic pathway namely GM2 synthase and GM1 synthase were increased in confluent cells compared to those grown under sparse conditions (Fig.7C). Interestingly the enzymes belonging to the early part of the SL biosynthetic pathway, namely SMS1, GCS or LCS were not altered (Fig.7C) suggesting that the expression of key enzymes determining the proportion between SM and GSLs (Fig.6C) was not altered under these conditions. Since there is no change in the expression levels of these key enzymes, we next examined whether a change in localization of enzymes could potentially underlie the change in GSL production. To this end, we studied the effect of addition of Brefeldin A that destroys enzyme organization **(Fig. S2)** on SL biosynthesis under both sparse and confluent conditions. We found that addition of Brefeldin A neutralizes the difference between sparse and confluent conditions with respect to GSL production (Fig.7D), suggesting that the observed difference in GSL biosynthesis between sparse and confluent conditions is likely due to an altered localization of SL biosynthetic enzyme(s). So, we examined the localization of GCS, the key enzyme that controls GSL production (Fig.6C) in sparse and confluent conditions. We found that GCS was mostly localized to the *trans*-Golgi in confluent conditions (Fig.7E) similar to what was observed earlier (Fig.3), when cells were cultured in near confluent conditions (see Methods). On the contrary, in cells grown in sparse conditions, the peak localization of GCS was in the *cis*-side of the Golgi (Fig.7E). These experiments together suggest that there is a difference in the localization of GCS between cells grown in sparse and confluence conditions and this likely accounts for the differences in SL biosynthesis in these cells. Then, we examined if GRASP55 contributes to this difference in localization of GCS and the subsequent change in SL biosynthesis between sparse and confluent fibroblasts. To this end, we first compared the localization of GCS in GRASP55 KO fibroblasts grown in sparse and confluent conditions. We found that in both these conditions, GCS was localized to the *cis*-side of the Golgi in GRASP55 KO fibroblasts (Fig.7E) similar to what was observed earlier in HeLa cells (Fig.3). Next, we examined if there is a confluence-induced alteration in SL biosynthesis in GRASP55 KO fibroblasts. As noted earlier, GRASP55 KO fibroblasts produced more GSLs compared to control cells, **(Fig. S5C)** and these increased levels of GSLs were not reduced when cells reached complete confluence (Fig.7F). This suggests that confluence associated reduction in GSL biosynthesis depends on GRASP55. These series of experiments suggest that when fibroblasts grow to reach confluence there is a GRASP55-dependent change in the localization of GCS from *cis*- to *trans*-Golgi and an associated reduction in GSL production.

**Figure 7.**
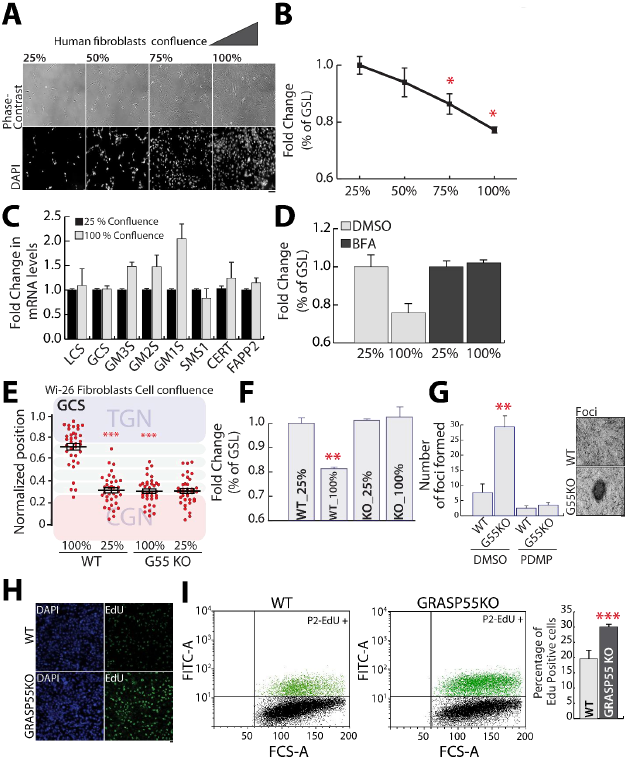
GRASP55 contributes to confluency-induced changes in GSL production: **(A)** WI-26 human fibroblasts grown to different levels of confluence stained with DAPI and imaged in phase contrast and fluorescence mode. Bar: 500μm. **(B)** Fibroblasts were grown to indicated confluency and SL output monitored by the [^3^H] - sphingosine pulse-chase assay. Total GSL levels were quantified and are represented as fold changes with respect to lowest confluence (25%). **(C)** Fibroblasts were grown to either complete or sparse (25%) confluence and mRNAs were extracted. The indicated mRNAs were quantified by RT-PCR and expressed as levels relative to those in cells grown in sparse condition. **(D)** Fibroblasts were grown to either complete or sparse (25%) confluence, treated with DMSO or Brefeldin A (BFA; 5μg/ml) for 30min and SL production monitored by the [^3^H] - sphingosine pulse-chase assay in the continued presence of DMSO or BFA for 8h. Total GSL levels were quantified and are represented as fold changes respect to control cells at low confluence (25%). **(E)** Fibroblasts grown to complete or sparse (25%) confluence, were transfected with GCS-HA and subsequently treated with nocodazole (33 μM) for 3 hours and then fixed and stained. Intra-Golgi localization of GCS was quantified by line scan analysis as described in Figure 3A-B. **(F)** GRASP55 KO fibroblasts were grown to complete or sparse (25%) confluence and SL production monitored by the [^3^H] - sphingosine pulse-chase assay. Total GSL levels were quantified and are represented as fold changes with respect to lowest confluence (25%). **(G)** Fibroblasts (WT and GRASP55 KO) were grown to confluence in 6 well plates and then maintained for 7 days in the presence or absence of PDMP (2 μM) to monitor foci formation. The cells were stained with crystal violet (0.25%) and the number colonies per well were counted and represented. Values are mean ± SD. Data representative of 2 independent experiments. **(H, I)** Control and GRASP55 KO fibroblasts were cultured same as in (A) then incubated with EdU overnight and either processed for FACS analysis (graph) or stained with DAPI (image). The number of EdU positive cells were quantitated by FACS and plotted. Flow cytometry distributions are representative of at least 3 independent experiments. Values are Mean ± SD. *p <0.05, **p <0.01, ***p <0.001 (Student’s t test).

Next, we explored if this GRASP55-mediated regulation of GSL biosynthesis that occurs when cells reach confluence translates into observable cell behaviour. When cells reached complete confluence under *in vitro* culture conditions the phenomenon of contact inhibition of growth is activated that prevents cell division. First, we examined the continuation of cell growth after reaching confluence by studying EdU incorporation (*55*). Control and GRASP55 KO fibroblasts were allowed to reach confluence by culturing for 7 days and the incubated with EdU for 3h. The number of cells incorporating EdU a marker for DNA replication was monitored by FACS analysis. We found that incorporation of EdU was at least 50% higher in GRASP55 KO fibroblasts compared to control fibroblasts (Fig.7H,I). Next, we examined if the absence of GRASP55 impaired contact inhibition of growth using focus-forming assay (*56*). Cells were cultured to complete confluence and maintained for 7-8 days before examining the number of foci formed. Control cells formed 4-5 foci per well, while GRASP55 KO cells had 4-fold more foci with 20-25 foci observed per well (Fig.7G). To understand if this difference is due to altered GSL biosynthesis we studied the effect of inhibition of GSL biosynthesis on foci formation. The difference in number of foci was ablated upon treatment with 1-phenyl-2-decanoylamino-3-morpholino-1-propanol (PDMP) a known inhibitor of GSL biosynthesis (Fig.7G). Thus, GRASP55-mediated reduction in GSL production associated with increased cell density contributes to contact inhibition of growth. Of note, increased GCS activity was correlated with tumour development in a mouse model of hepatocellular carcinoma (*57*) and mouse insertional mutagenesis studies have found that inactivation of GRASP55 promotes tumour transformation and/or growth in liver and colorectal cancer models (Ref: Candidate Cancer Gene Database). In light of these observations, regulation of contact inhibition of growth by GRASP55-mediated alteration of GSL biosynthesis may have potential pathological relevance.

## Discussion

Cell surface glycans play important roles in cell-cell interaction and are synthesized mainly by the Golgi apparatus. The glycosylation enzymes that assemble these glycans are compartmentalized to one or more cisternae of the Golgi stack. The molecular mechanisms of Golgi compartmentalization and its contribution to regulation of glycosylation have remained unclear. Here we describe, for the first time, a regulator of Golgi enzyme compartmentalization - GRASP55 that compartmentalizes two key GSL biosynthetic enzymes – GCS and LCS. It acts by binding to these enzymes, preventing their entry into retrograde transport carriers and thus retaining them in the cisterna. This *retainer* action of GRASP55 opposes COPI (and GOLPH3) mediated retrograde transport of residents and a balance between these two actions compartmentalizes the enzymes to trans-Golgi. This enzyme compartmentalization controls cargo flux between competing SL biosynthetic reactions by providing privileged access of enzymes to the cargoes and thus controls sphingolipids produced by the Golgi. GRASP55 thus controls cell surface sphingolipids profile through enzyme compartmentalization and in turn regulates density-dependent cell growth. We propose that two major cellular processes act together to define the glycan profile of a cell - transcriptional processes that regulate the expression of glycosylation factors to determine the type of glycans that will be produced, and membrane transport processes in the Golgi regulate the levels (*23*) and localization of enzymes to determine the proportion of each glycoform produced. We discuss below the implications of our study for the cell biology of Golgi organization as well as glycobiology.

### Mechanisms of bidirectional transport across the Golgi

Transport across the Golgi apparatus is bidirectional with forward or anterograde transport happening from *cis* to *trans*-Golgi and retrograde transport in the opposite direction. According to the cisternal progression/maturation model, the anterograde flux is mediated by cisternal progression and the retrograde flux by COPI mediated processes (*16, 17*). The mechanistic basis of retrograde transport is relatively well established with recent studies identifying specificity encoders in this transport process (*19, 23, 27, 28*). The cisternal progression, on the other hand, is considered a bulk-flow/passive process with the retention of proteins in the progressing cisterna secondary to their inefficient interaction with the COPI machinery. Surprisingly, we discovered that GRASP55 actively prevents the incorporation of proteins into retrograde COPI vesicles by binding and *retaining* them in the maturing /progressing cisterna. This action of GRASP55 favours the anterograde progression of its client molecules and suggests that anterograde transport by cisternal progression is not a passive but an actively regulated process. Further, recent studies have shown that at least some secretory cargoes have a tendency to enter retrograde transport pathways (*58*) and therefore may require a “retainer” like action to promote their efficient anterograde transport.

### Mechanisms regulating Golgi compartmentalization

The retention of glycosylation enzymes in the Golgi apparatus, is due to their continuous retrograde transport by the COPI machinery. How the same retrograde transport machinery achieves the compartmentalization of enzymes to different cisterna was not clear. The discovery of an enzyme adaptor – GOLPH3 acting on the trans-side of the Golgi, suggested that the *trans* and *cis*-Golgi localized enzymes may have differential modes of interacting with the COPI machinery. While the cis-Golgi localized proteins interacted with COPI directly (*19*), the trans-Golgi residents may require an adaptor recruited specifically to the trans-Golgi, thus allowing the same COPI machinery to compartmentalize enzymes to two different sub-Golgi compartments. Our discovery of GRASP55 as an active retainer shows this process is more elaborate and with both retaining and recycling adaptors required for appropriate localization to the Golgi. Indeed, in the absence of GRASP55 and with only GOLPH3 present, the compartmentalization of LCS is altered. Thus, compartmentalization of Golgi residents is a dynamic process resulting from a balance between their anterograde and retrograde flux mediated by retaining and recycling adaptors respectively (Fig.5G). Recently, GRASP proteins have been suggested to inhibit COPI vesicle formation (*43, 59*). This activity of GRASP55 may further contribute to oppose the retrograde transport of residents. Given the well-known regulation of GRASP55 and GOLPH3 activities by phosphorylation in response to various stimuli, we expect that regulation of glycosylation by altering Golgi compartmentalization may turn out to be a regulated physiological phenomenon.

We also note that there are several other Golgi resident proteins with canonical PDZ domain binding motifs, which includes other enzymes and channels. Thus, GRASP55 (and possibly GRASP65) may play a wider role in the localization of several proteins in the Golgi. Further, GRASP55 as a prototype “retainer”, provides insights to identify similar molecules. We surmise that a retainer molecule: a. will be a peripheral membrane protein dynamically associated to the Golgi [else it will lead to the conundrum of who retains the retainer? (Elko. personal communication)]; b. binds to Golgi residents; and c. is excluded from the peri-Golgi vesicles. There are several such molecules in the Golgi including other Golgins and thus we are optimistic that other “retainers” will be found.

### Contribution of Golgi to fidelity of glycan biosynthesis

How and to what extent does Golgi play an active role in controlling glycan output remained unclear. The ordered arrangement of sequentially acting enzymes of N-glycan biosynthesis in Golgi implied that their localization might contribute to glycan output (*29*). Nonetheless, experiments have shown that a single compartment could still support sequential N-glycan processing (*60–63*). In retrospect, compartmentalization for sequentially acting enzymes seems superfluous, nevertheless it may have some role (*30*). Here, we show that Golgi plays a substantive role in regulating the glycan output especially in pathways involved competing reactions., such that when Golgi organization is destroyed there is nearly 2-3 fold change in glycan output (Fig.1E,F). These data and those presented by others (*64, 65*) clearly indicate that enzyme localization determines output of competing reactions. When two enzymes compete for a substrate, a privileged access to it (by differential positioning along the pathway) can increase the synthesis of glycans produced by the enzyme. While different from what is presented here, previous studies on metabolic bias in sphingosine processing in lysosome *vs* mitochondria (*66*) and by Golgi to ER translocation of O-glycan initiating enzymes (*67*) suggest that localization-dependent regulation of glycosylation, and metabolism in general, may be a wide-spread phenomenon.

### Regulation of glycosylation by GRASP55

GRASP55 has been suggested to play an important role in the stacking of cisternae (*41, 42*), formation of Golgi ribbon (*68*), conventional as well as unconventional secretion (*45–47*) and glycosylation (*43, 48*). It has been proposed to regulate glycosylation by regulating the kinetics of intra-Golgi transport (*43*) and/or ribbon formation (*48*). For GRASP55 mediated regulation of GSL biosynthesis, neither is a likely explanation since kinetics of GSL biosynthesis **(Fig. S5A)** is very different from that of protein glycosylation and downregulation of several matrix proteins known to fragment Golgi ribbon do not affect GSL biosynthesis (Fig.2A). Instead, we show that absence of GRASP55 leads to a change in the localization of LCS and GCS (Fig.3). This alters SL production by providing a privileged early access of substrates to these enzymes and/or providing them a more permissive environment (*69, 70*) and thus increases GSL production (Fig.6E). Thus, our study here provides an alternate or complementary view of how GRASP55 regulates glycosylation.

### SL biosynthesis

SL biosynthesis depends on CERT mediated transfer of ceramide from ER to TGN for SM biosynthesis (*71*) and FAPP2-mediated transfer of GlcCer from *cis*-Golgi to TGN for Gb3 biosynthesis (*34*). Earlier reports had shown that GCS and SMS1 are present in *cis*- and *trans*-Golgi respectively (*35*). Surprisingly, we observe that GCS is present in the *medial/trans*-Golgi. We ascribe the observed difference in localization of GCS to the use of constructs with a tag that blocks the C-terminus (important for interaction with PDZ domain of GRASP55) in the previous studies (*35*). This revised localization of GCS has implications for the model of SL biosynthesis: if GCS is localized to *trans*-side why is the activity of GCS not sensitive to CERT KD? While GCS is indeed present in the *trans*-Golgi, a large fraction of SMS1 is present in a distinct compartment, the TGN. CERT-mediated transfer of ceramide likely happens at the TGN and thus SMS1 depends on ceramide delivered by CERT unlike GCS.

To sum up, we identify a novel molecular pathway regulated by GRASP55 that compartmentalizes specific glycosylation enzymes and by this action modulates the competition between reactions to achieve a specific cellular glycan profile.

## Acknowledgments

We thank Antonella De Matteis for valuable discussions, Nina Dathan for help with cloning and construct preparation and Francesco Russo for assistance with statistical analysis. We thank the Bioimaging Facility of the Institute of Biochemistry and Cell Biology and the joint FACS Facility of the Institute of Genetics and Biophysics (IGB-CNR, Naples) and IBBC-CNR for access to technologies and Pasquale Barba for assistance with FACS. We thank Dr. Bruno Correia, EPFL, Switzerland, for help with the analysis of protein-protein interaction studies. We thank Drs. Corda, Colanzi, Jung, Grimaldi, Roy, Qadri, Krishnamoorthy, Ahamarshan and Polishchuk for critical comments. We thank MEDINTECH, the Italian Node of Euro-Bioimaging (Preparatory Phase II – INFRADEV), Italian Cystic Fibrosis Research Foundation (FFC #6-2019), POR Campania project 2014-2020 (C.I.R.O.) and CTC project for financial support. R.R. acknowledges financial support from Fondazione Italiana per la Ricerca sul Cancro (FIRC Fellowship 15111). A.L. acknowledges financial support from the, AIRC (Projects IG 15767 and IG 20786), TERABIO, PON-IMPARA and S.A.T.I.N. G.D.A. acknowledges financial support from EPFL institutional fund, Kristian Gerhard Jebsen Foundation and from the Swiss National Science Foundation (SNSF) (grant number, 310030_184926).

## Author contributions

PP contributed to development of the idea, conducted most of the experiments, analysis and interpretation of data and contributed to writing the manuscript; IA conducted the experiments on peptide binding using purified proteins; MP was involved in labelling and acquisition of EM and confocal microscopy images; RR assisted in acquisition and analyses of electron micrographs; DR conducted mRNA analysis; GT was involved in EM sample preparation and acquisition of images; LC conducted MALDI-MS and lipidomics measurements; CJCG performed the ITC experiments; GV assisted PP in morphological analysis of Golgi; ND was involved in cloning and construct preparation; JSY performed vesicle budding assay and was guided in this by VWH; PH was involved in synthesis of biotinylated peptides; JN provided the WT and GRASP55 KO fibroblasts, performed the protein-protein interaction studies and was guided by MP; TN, JI, MJH, LO and YH contributed to MS analysis of lipids; AB performed *in silico* modelling of GRASP55-GCS peptide interaction and was guided by MT; AL contributed to development of the idea of how Golgi organization may play role in regulating glycosylation and the mode of action of GRASP55 in regulating enzyme compartmentalization, critical analysis of the data throughout the entire project and also provided inputs in manuscript preparation. GDA contributed to setting up the system to monitor SL biosynthesis, provided important inputs on the data to understand and interpret changes in lipid biosynthesis, and also provided critical inputs in manuscript preparation. SP developed the idea, designed and supervised the entire project, and wrote the manuscript.

