## Supplementary material for "Regulated compartmentalization of enzymes in Golgi by GRASP55 controls cellular glycosphingolipid profile and function": Supplementary Information.pdf

Supplementary Information contains:

1. Supplementary Figures and corresponding legends (**S1-S10)**

2. Methodology

3. Supplementary Tables (**S1-S8**)

4. Supplementary Bibliography

### Figure S1

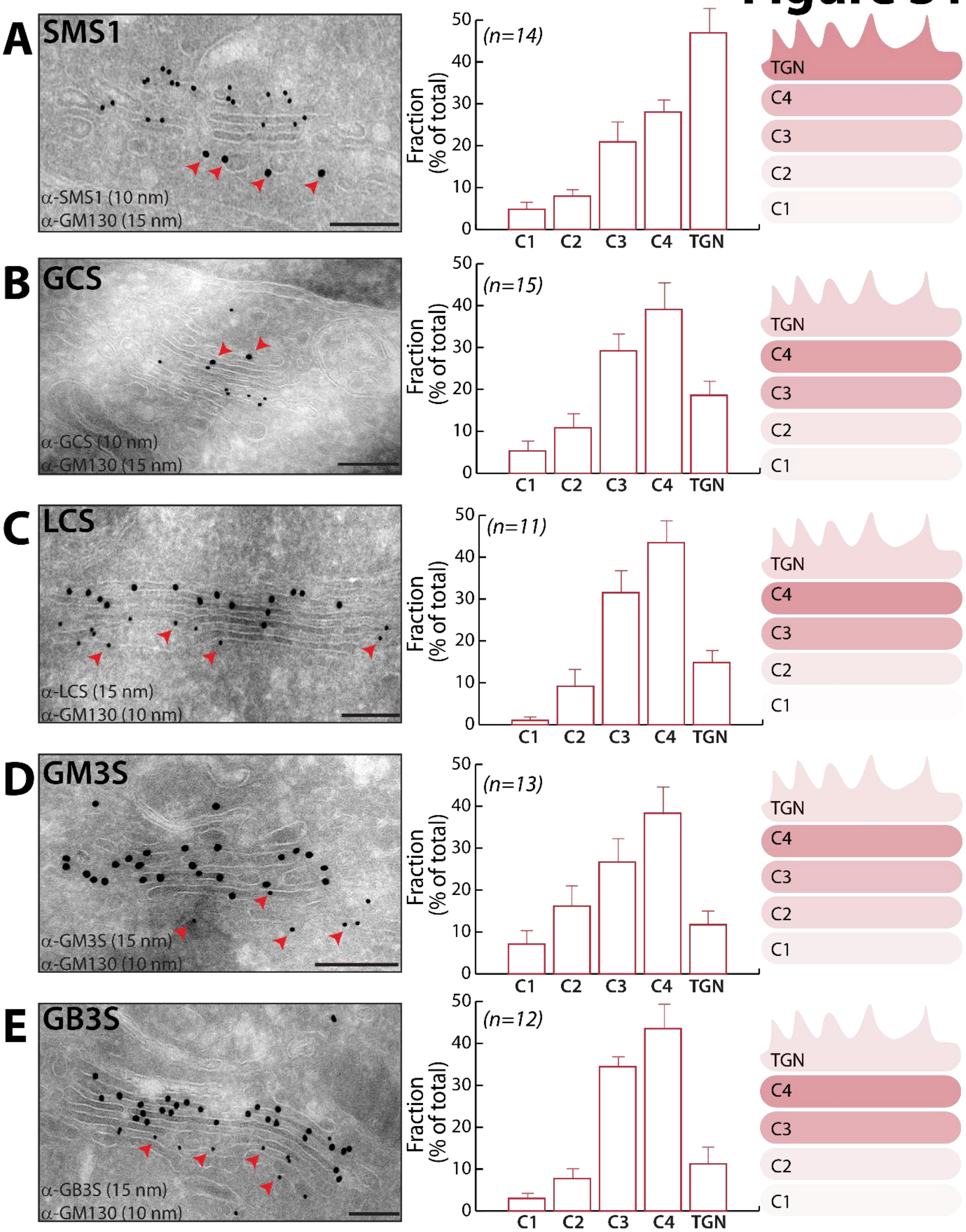
A
SMS1
α-SMS1 (10 nm)
α-GM130 (15 nm)
(n=14)
Fraction (% of total)
C1 C2 C3 C4 TGN
TGN
C4
C3
C2
C1
B
GCS
α-GCS (10 nm)
α-GM130 (15 nm)
(n=15)
C
LCS
α-LCS (15 nm)
α-GM130 (10 nm)
(n=11)
D
GM3S
α-GM3S (15 nm)
α-GM130 (10 nm)
(n=13)
E
GB3S
α-GB3S (15 nm)
α-GM130 (10 nm)
(n=12)

**Figure S1. Distribution of SL biosynthetic enzymes in Golgi:**

**(A-E)** HeLa cells were transfected with indicated HA-tagged SL biosynthetic enzymes for 16 hours, fixed and processed for cryoimmunolabeling with anti-HA and anti-GM130 antibodies followed by Protein A-gold. HA and GM 130 are represented by 10- and 15-nm gold particles in case of SMS1 and GCS while in case of LCS, GB3S and GM3S they correspond to 15- and 10-nm gold particles respectively. Red arrow heads indicate cis face of Golgi marked by GM130 labelling. Enzyme distribution expressed as fraction of total gold particles per Golgi stack including TGN. (n indicated in the graph); data are mean ± SEM, Scale Bar, 200nm.

### Figure S2

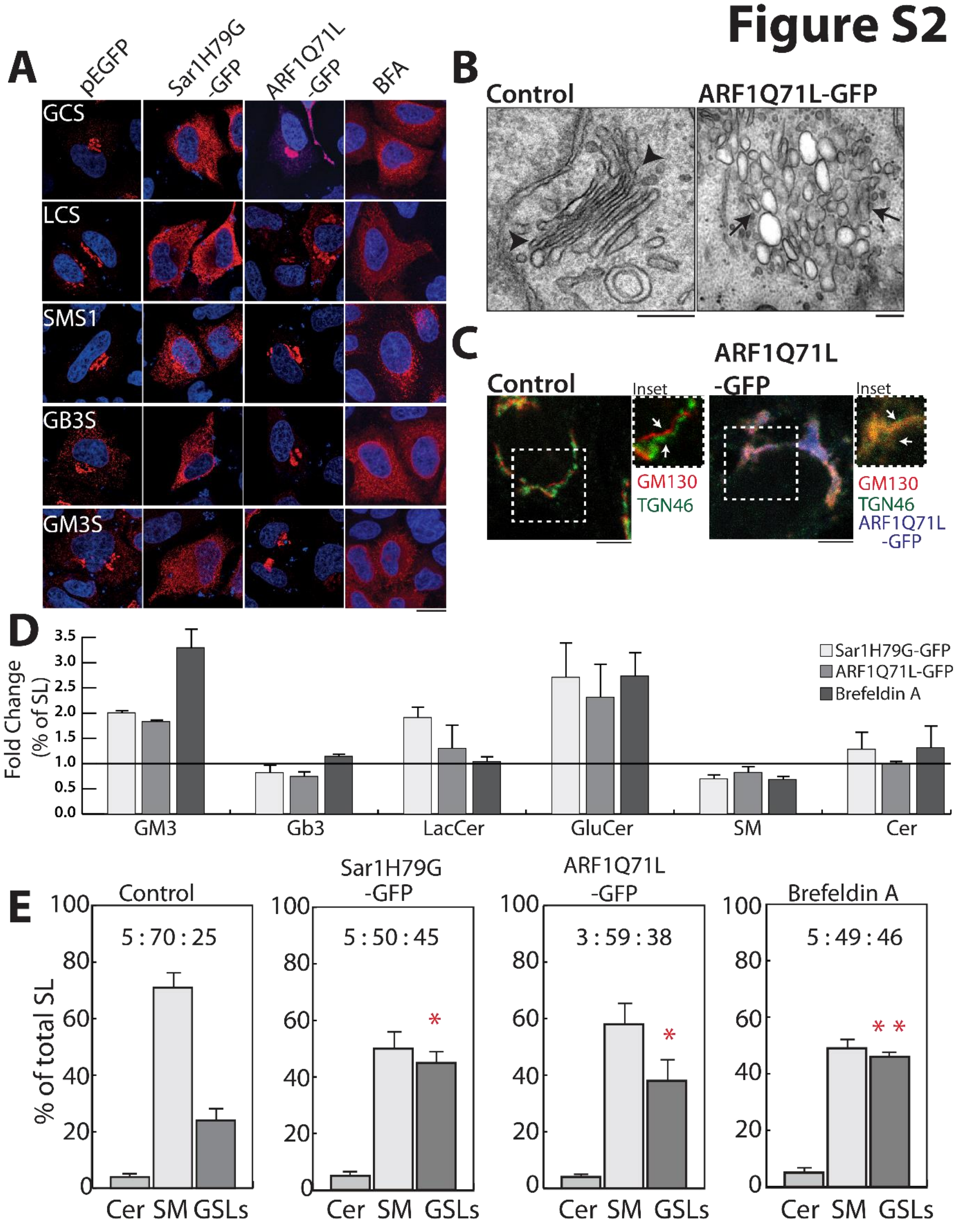
A
pEGFP
Sar1H79G -GFP
ARF1Q71L -GFP
BFA
GCS
LCS
SMS1
GB3S
GM3S
B
Control
ARF1Q71L-GFP
C
Control
Inset
ARF1Q71L -GFP
Inset
GM130
TGN46
GM130
TGN46
ARF1Q71L -GFP
D
Fold Change (% of SL)
3.5
3.0
2.5
2.0
1.5
1.0
0.5
0.0
GM3
Gb3
LacCer
GluCer
SM
Cer
Sar1H79G-GFP
ARF1Q71L-GFP
Brefeldin A
E
% of total SL
Control
5 : 70 : 25
Sar1H79G -GFP
5 : 50 : 45
ARF1Q71L -GFP
3 : 59 : 38
Brefeldin A
5 : 49 : 46
100
80
60
40
20
0
Cer SM GSLs
*
* *

**<u>Figure S2. Golgi organization determines faithful SL output:</u>**

**(A)** HeLa cells were transfected with indicated HA-tagged SL biosynthetic enzymes and either empty vector (PEGFP) or a plasmid encoding Sar1H79G-GFP or ARF1Q71L-GFP for 16 hours or treated with brefeldin A (BFA) (5µg/ml) for 30 min, fixed, permeabilized and stained with DAPI (blue) and anti-HA antibody (red). Scale bar, 10 µm. **(B)** HeLa cells transfected with PEGFP or ARF1Q71L-GFP for 16 hours were fixed and processed for electron microscopy. Black arrow heads indicate the intact Golgi and black arrows represent the tubulo-vesicular clusters. Scale Bar, 200nm. **(C)** HeLa cells were transfected with ARF1Q71L-GFP for 16 hours, fixed, permeabilized and stained with anti-GM130 and anti-TGN46 antibodies. GM130 is represented in red, TGN46 in green, and GFP in blue. White arrows indicate the separation of cis and trans markers of the Golgi in control cells and their overlap in ARF1Q71L-GFP expressing cells. Scale bar, 10 µm. **(D-E)** SL species quantified by radioactive pulse chase assay in HeLa cells transfected with Sar1H79G-GFP or ARF1Q71L-GFP or treated with BFA and represented as fold change with respect to control (**D**) or as relative percentage of Cer, SM and GSLs **(E)**. For BFA treated cells, the SL output was measured 8h after a pulse. Data represented as mean ± SD of 2 independent experiments. *$P < 0.05$, **$P < 0.01$ (Student's t test).

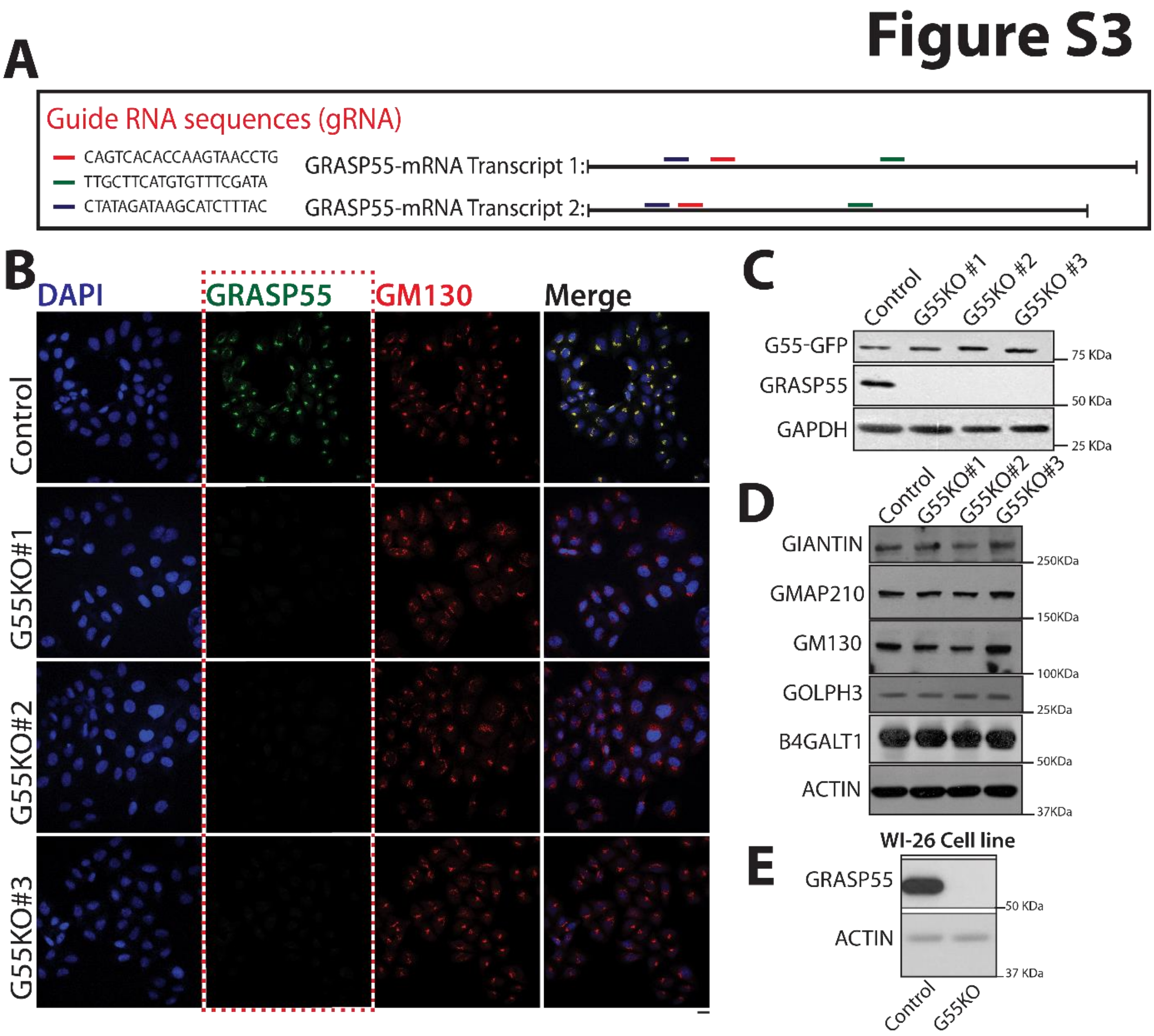


**<u>Figure S3. Generation of GRASP55KO cell lines:</u>**

**(A)** Specific guide RNA oligonucleotides for human GRASP55 gRNA #1, #2 and #3 used for CRISPR/Cas9-mediated deletion of GRASP55 expression. The sequences of oligonucleotides are shown and the sites in GRASP55 transcripts to which the gRNA correspond are indicated. **(B)** The control and GRASP55 KO clones were fixed, permeabilized and stained with DAPI (blue), anti-GM130 antibody (red) and anti-GRASP55 antibody (green). Scale bar, 10 µm. **(C)** Control and GRASP55 KO clones transfected with GRASP55-GFP were analysed by Western blotting with anti-GRASP55 and anti-GFP antibody. **(D)** Cell lysates from control and GRASP55 KO clones were analysed for the expression of indicated Golgi-Matrix proteins by western blotting. **(E)** Cell lysates of wild type and GRASP55 KO fibroblasts were analysed by western blotting for GRASP55 expression.

### Figure S4

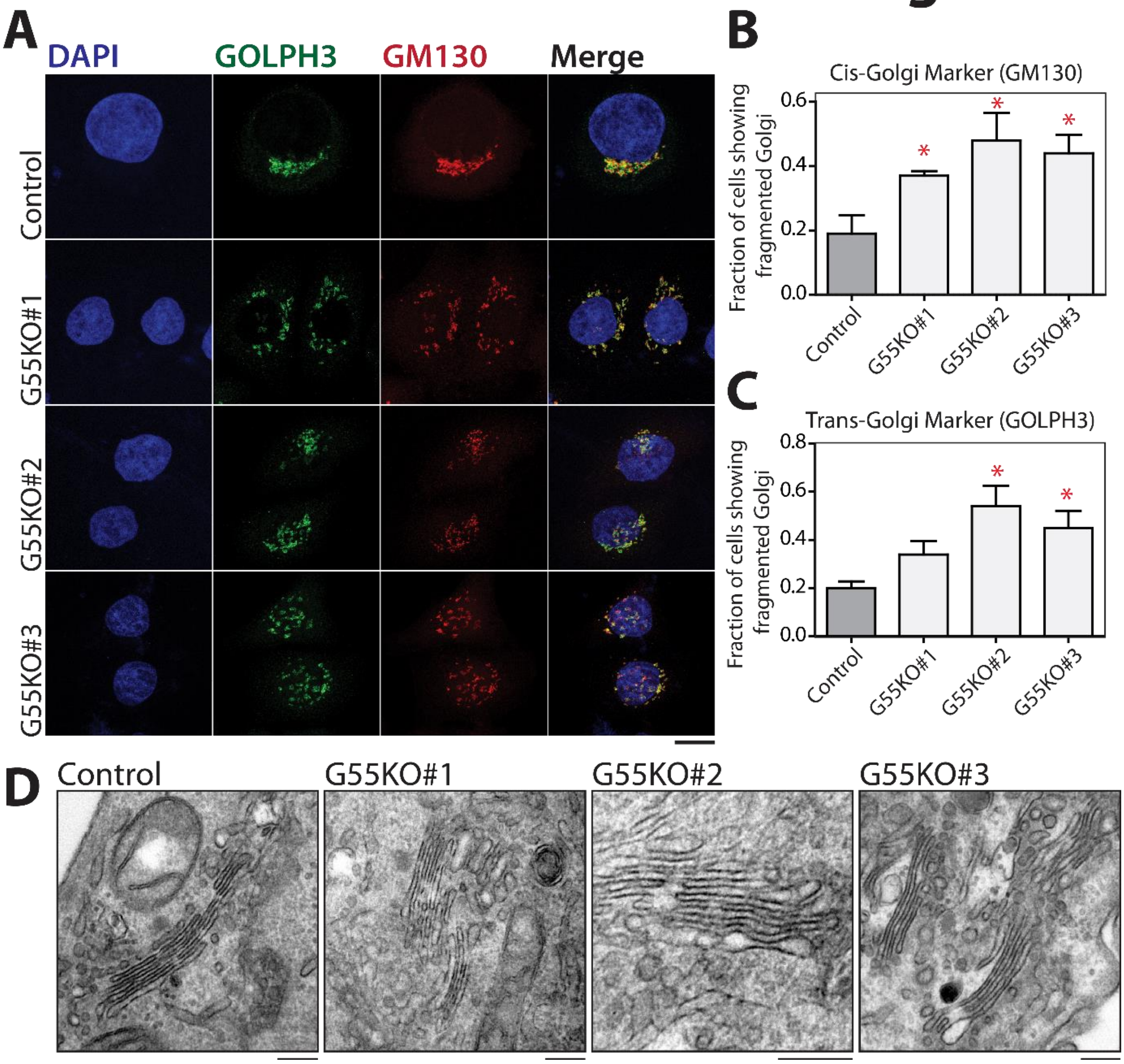


**Figure S4. Characterization of Grasp55 KO clones:**

**(A)** Control and GRASP55 KO clones were fixed, permeabilized and stained with DAPI (blue), anti-GM130 antibody (red) and anti-GOLPH3 antibody (green). The images represent the maximum intensity projection along the z-axis of 7 consecutive confocal sections. Scale bar, 10 µm. **(B-C)** The graphs represent the fraction of cells displaying fragmented Golgi in control and GRASP55 KO clones. Values are mean ± SD of 2 independent experiments. *p <0.05 (Student's t test). **(D)** Control and GRASP55 KO cells were fixed and prepared for EM. The Golgi profile shown suggests no significant alterations in the stack architecture of Golgi. Scale bar corresponds to 200nm.

### Figure S5

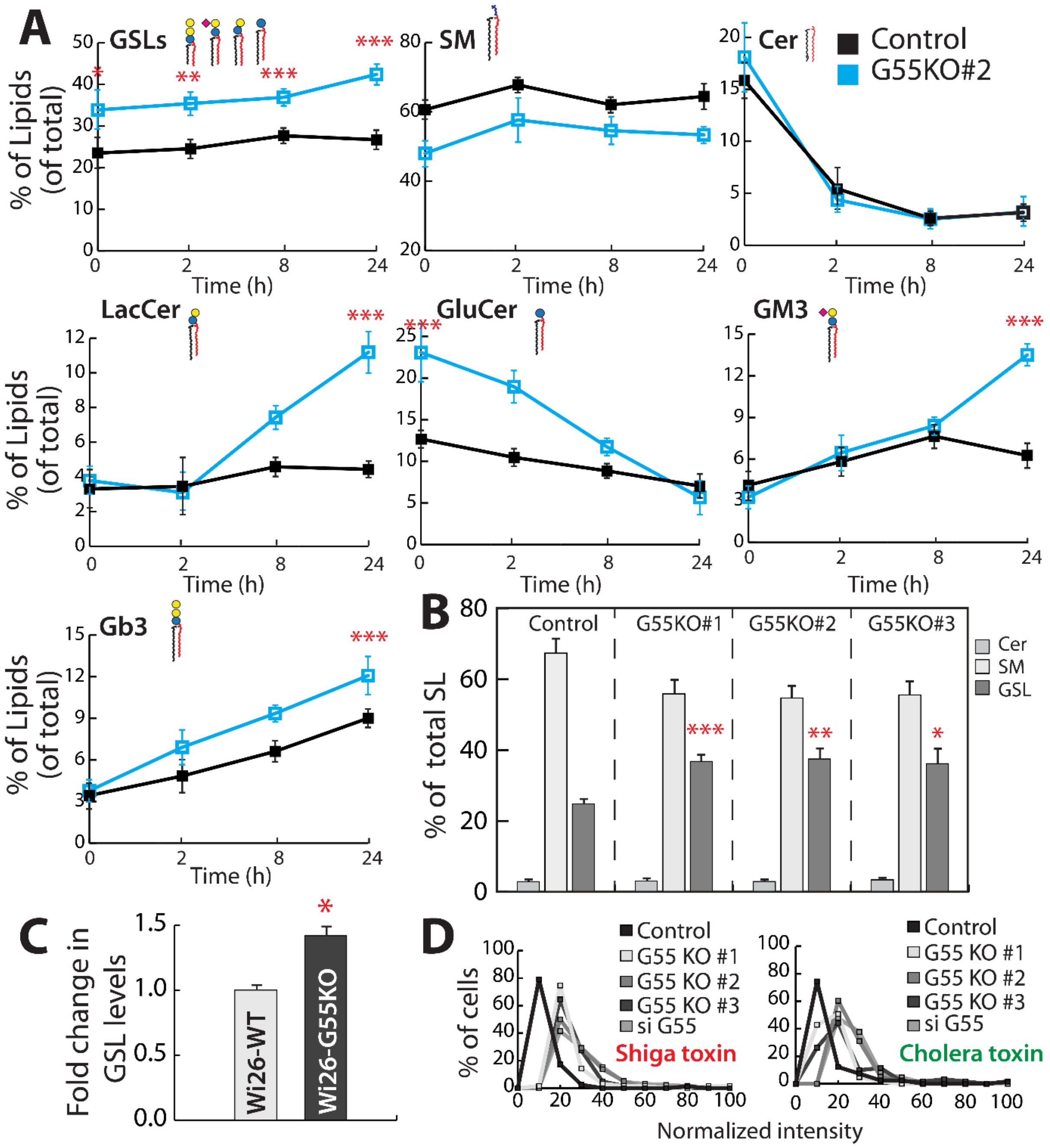


**Figure S5. GRASP55 regulates GSL biosynthesis:**

**(A)** Control and GRASP55 KO (#2) cells were subjected to radioactive pulse chase assay as described earlier, the indicated SL species were quantified and are represented as percentage of total SLs. X-axis represents chase time. Data represented as mean ± SD of 3 independent experiments **(B)** Control and GRASP55 KO cells were subjected to radioactive pulse chase assay as described earlier. SL levels were quantified after 24h of chase and represented as percentage of total SLs.

Data represented as mean ± SD of 3 independent experiments. **(C)** SL production in control or GRASP55 KO fibroblasts was measured by [$^3$H] - sphingosine pulse-chase assay, and total GSL levels are expressed as fold changes with respect to control. Data represented as mean ± SD of 2 independent experiments. *p <0.05, **p <0.01, ***p <0.001 (Student's t test). **(D)** Effect of GRASP55 depletion on GSL levels measured by Cy3-conjugated ShTxB (Shiga Toxin) and Alexa488-conjugated ChTxB (Cholera Toxin) staining in control and GRASP 55 KO cells or cells treated with GRASP55 siRNA. The cells were imaged by epifluorescence microscopy and intensity of the fluorescence staining quantitated. Frequency distribution of level of STxB and CTxB staining is represented with intensities normalized to maximum intensity for each condition (>100 cells per condition were analysed).

### Figure S6

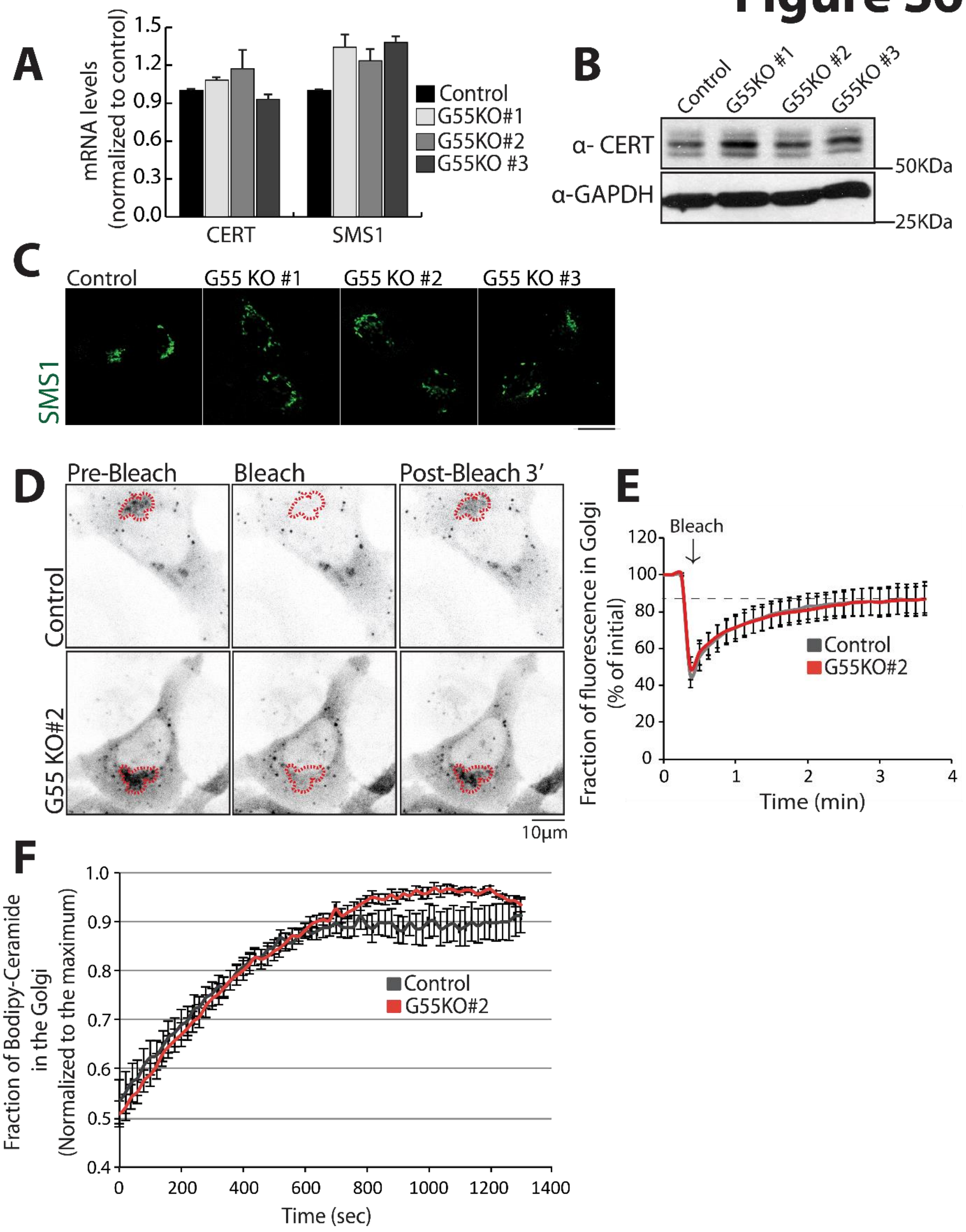
A
mRNA levels
(normalized to control)
1.5
1.2
0.9
0.6
0.3
0.0
CERT
SMS1
Control
G55KO#1
G55KO#2
G55KO #3
B
Control
G55KO #1
G55KO #2
G55KO #3
α- CERT
α-GAPDH
50KDa
25KDa
C
Control
G55 KO #1
G55 KO #2
G55 KO #3
SMS1
D
Pre-Bleach
Bleach
Post-Bleach 3'
Control
G55 KO#2
10μm
E
Fraction of fluorescence in Golgi
(% of initial)
Bleach
120
100
80
60
40
20
0
1
2
3
4
Time (min)
Control
G55KO#2
F
Fraction of Bodipy-Ceramide
in the Golgi
(Normalized to the maximum)
1.0
0.9
0.8
0.7
0.6
0.5
0.4
0
200
400
600
800
1000
1200
1400
Time (sec)
Control
G55KO#2

**Figure S6. GRASP55 does not regulate SM biosynthetic arm of SL biosynthetic pathway:**

**(A)** The mRNA levels of CERT and SMS1 were evaluated by qRT-PCR (values are mean ± SD; n=3). **(B)** Control and GRASP55 KO cell protein lysates were analysed by western blotting for CERT expression. **(C)** Control and GRASP55 KO cells expressing SMS1-HA were fixed, and processed for immunofluorescence with anti-HA antibody (green). Scale Bar, 10 µm. **(D-E)** Control cells and GRASP55 KO (#2) cells were transfected with CERT-GFP for 16 hours, the area of Golgi indicated by the red dotted line was bleached and the recovery of fluorescence was observed by live epifluorescence imaging. Representative images of indicated times are shown (**D**). The ratio of fluorescence of the bleached area to an adjacent unbleached area was measured for each time point, normalized to initial values and plotted in the indicated graph **(E)**. Scale bar, 10 µm. **(F)** Kinetics of Ceramide transport to Golgi was studied using BODIPY labelled-C6 ceramide in GRASP55 KO cells. Cells were labelled with BODIPY C6 ceramide (10 µM) for 30 min at 4ºC. The cells were then washed and shifted to 37ºC in the microscope and analysed by epifluorescence microscopy. The perinuclear concentration of fluorescence signal was quantified and plotted. *$p < 0.05$ (Student’s t test).

### Figure S7

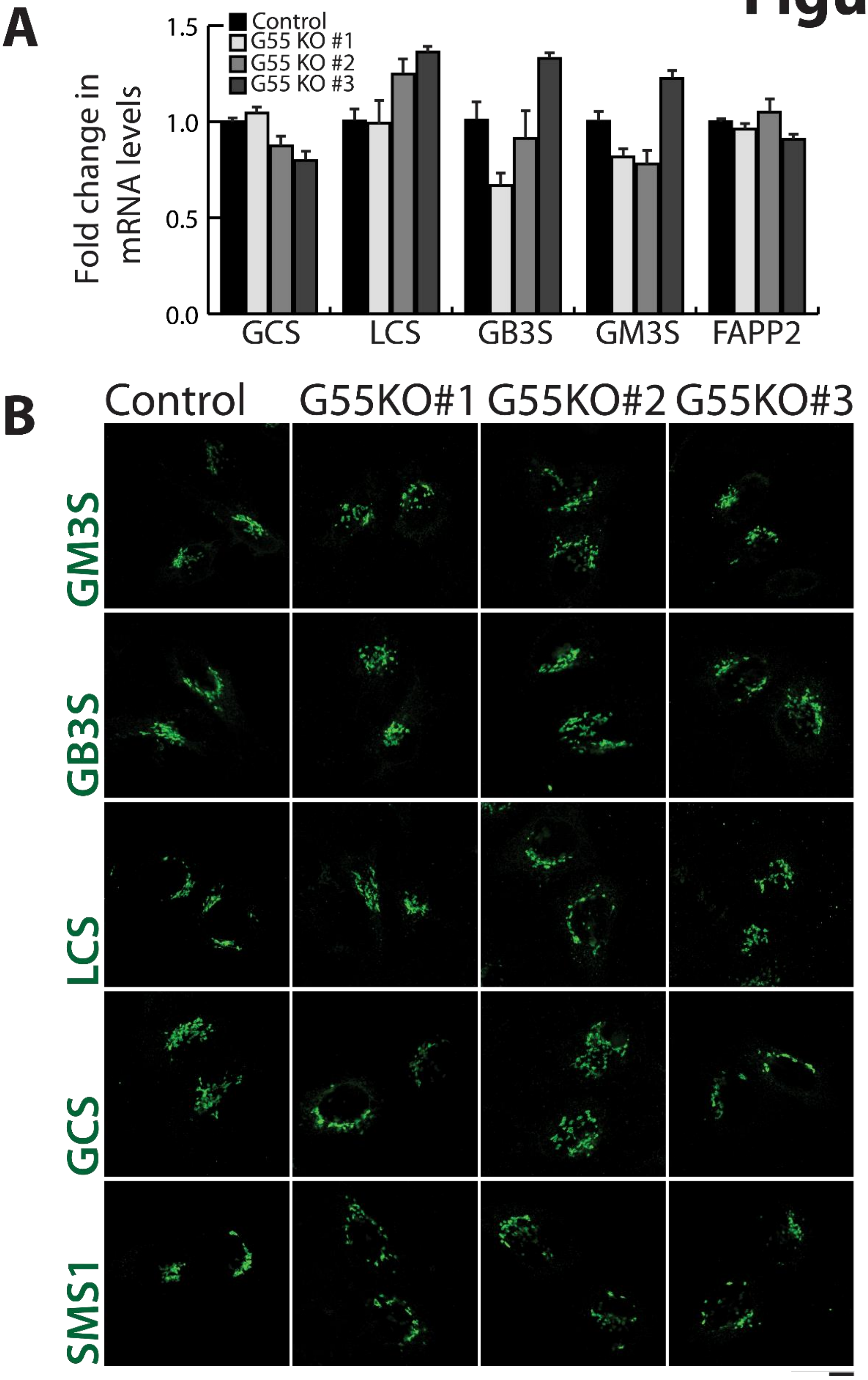

A
Control
G55 KO #1
G55 KO #2
G55 KO #3
Fold change in mRNA levels
1.5
1.0
0.5
0.0
GCS
LCS
GB3S
GM3S
FAPP2
B
Control
G55KO#1
G55KO#2
G55KO#3
GM3S
GB3S
LCS
GCS
SMS1

**Figure S7. GRASP55 does not regulate the levels or cellular localization of GSL biosynthetic enzymes:**

**(A)** Expression of GSL biosynthetic enzymes in control and GRASP55 KO clones were analysed by qRT-PCR. Values of mean ± SD. **(B)** Control and GRASP55 KO clones were transfected with the indicated HA-tagged GSL biosynthetic enzymes for 16 hours, fixed, permeabilized and stained with anti-HA antibody (Green). Scale bar, 10 µm.

### Figure S8

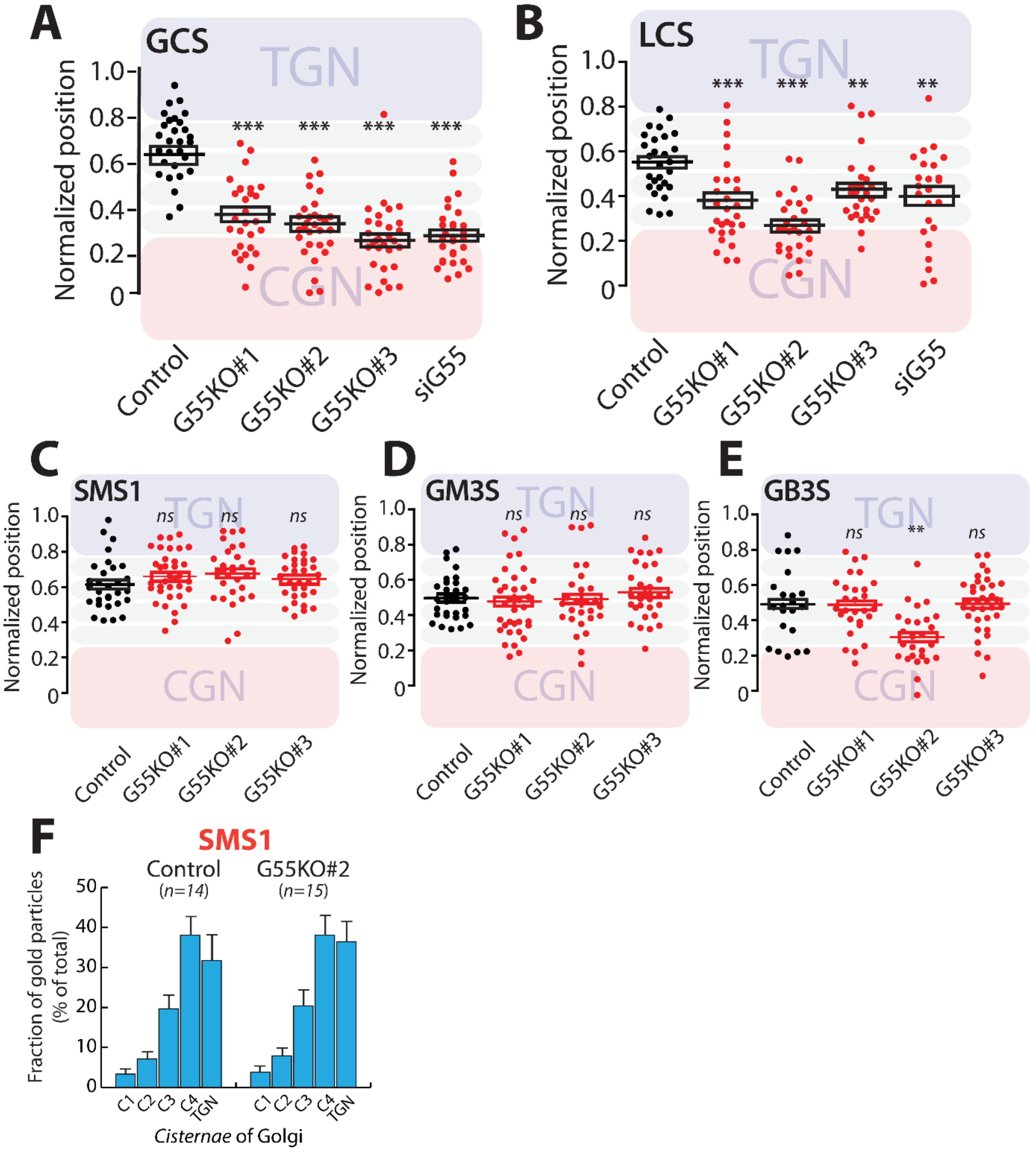


**Figure S8. GRASP55 regulates intra-Golgi localization of GCS and LCS:**

**(A-E)** Control cells treated with or without GRASP55 siRNA and GRASP55 KO clones were transfected with HA-tagged GSL biosynthetic enzymes, treated with nocodazole (33 µM) for 3 hours, and processed for immunofluorescence with anti-HA, anti-GM130, and anti-TGN46 antibodies. The relative position of HA-tagged enzymes with respect to GM130 and TGN46 was measured by line scanning and expressed as normalized positions of the peak intensity with peak of GM130 set to 0 and that of TGN46 to 1. The images are representative of >30 stacks analysed for each condition from three independent experiments. The data are mean ± SD from 3 independent experiments. **p <0.01, ***p <0.001(Student's t test) and *ns* signifies not statistically significant. **(F)** Control and GRASP55 KO cells were transfected for 16 hours with SMS1-HA and processed for cryoimmunolabeling. Distribution of indicated enzymes across the Golgi stack was quantified and represented as fraction of Gold particles in each cisterna (n indicated in the graph; data are Mean ± SEM).

### Figure S9

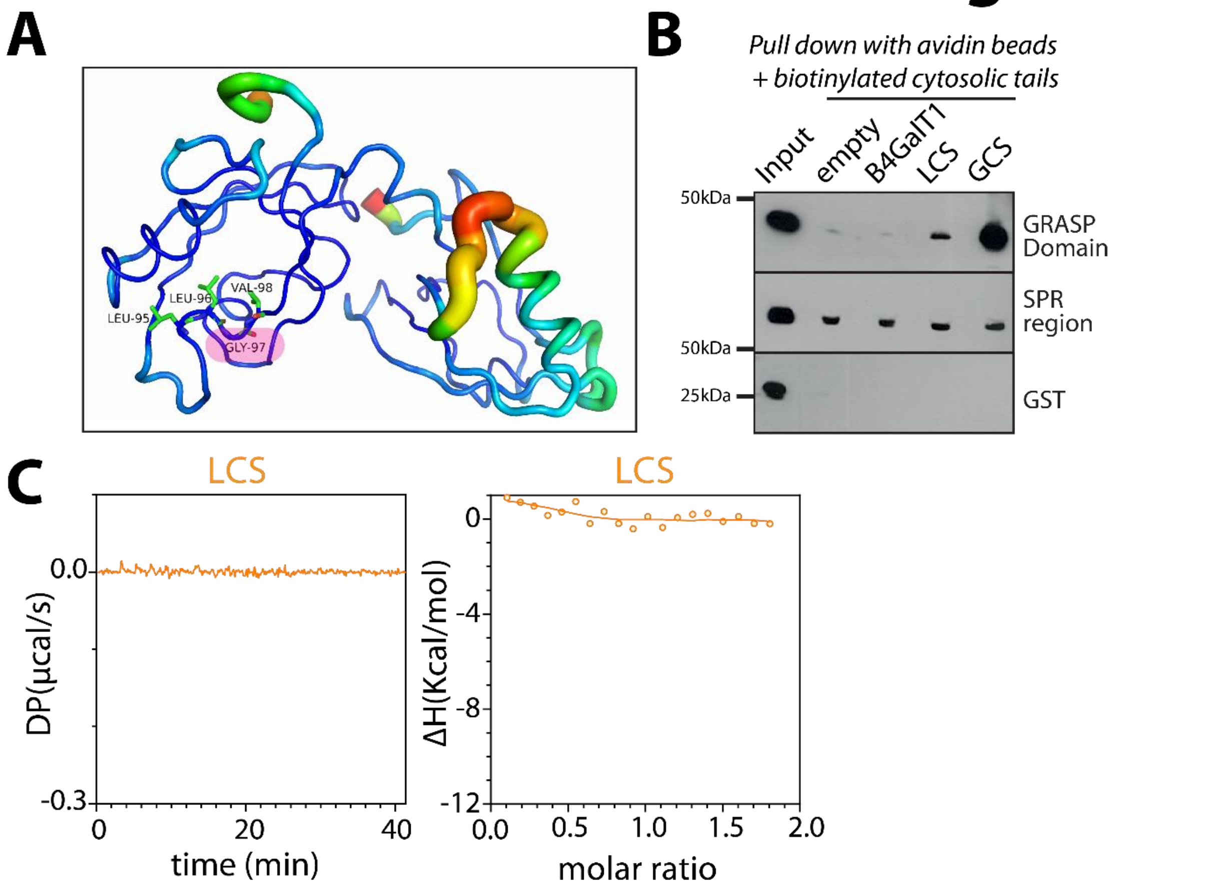


**Figure S9. GRASP55 directly interacts with GCS and LCS to promote their sub-Golgi localization:**

**(A)** B-factor analysis of the high-resolution crystal structure of complex GRASP55:Golgin45, revealed that the cognate carboxylate binding motif is rigid in nature. Glycine is more favoured such a 'strained conformation' (left handed alpha helical), where the loop is rigid. Asp is not a favoured residue to be in left-handed α helical conformation. It can be predicted, in the absence of mutant crystal structure, that there has been reorientation in the backbone conformation, which may have led to the disruption of the characteristic hydrogen bond network between the protein and peptide. **(B)** Chemically synthesized biotinylated peptides corresponding to cytosolic portions of glycosylation enzymes and indicated purified GST-tagged GRASP domain or SPR region of GRASP55 were incubated together and their interaction was monitored by pulling down the biotinylated peptides with avidin beads followed by western blotting with anti-GST tag antibody. **(C)** ITC profile, representative of at least

3 independent experiments, for biotinylated LCS cytosolic tails with recombinant GRASP55.

### Figure S10

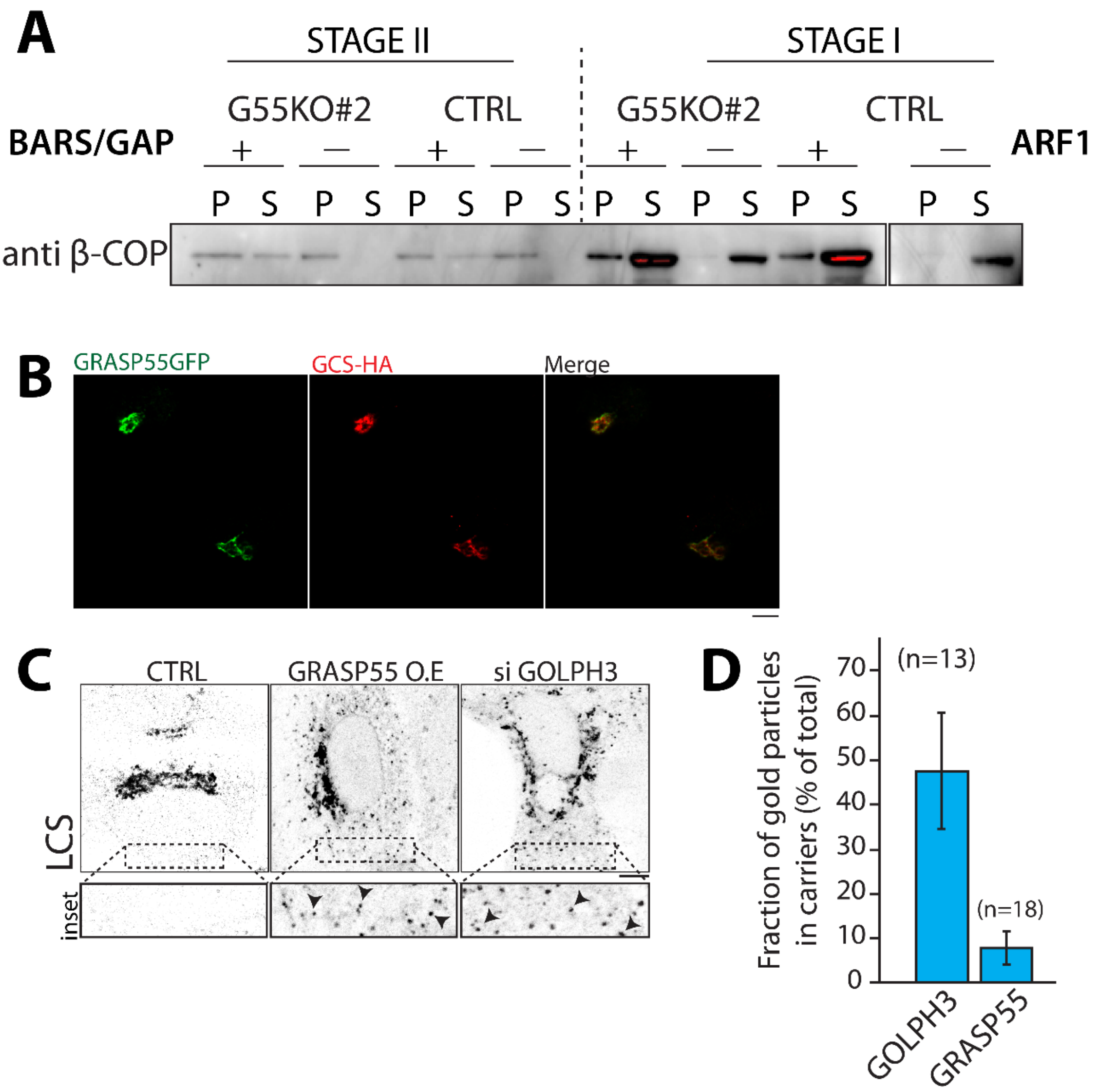


**<u>Figure S10. GRASP55 compartmentalizes the enzymes by preventing their entry into retrograde carriers:</u>**

**(A)** The two-stage incubation system was performed to reconstitute COPI vesicles. The first-stage incubation results in the ARF1-dependent recruitment of coatomer to Golgi membrane, as reflected by β-COP being redistribution from the supernatant (soluble) fraction to the pellet (Golgi membrane) fraction. The second-stage incubation results in the generation of COPI vesicles from Golgi membrane, as reflected by β-COP being redistributed from the pellet (Golgi membrane) fraction to the supernatant (vesicular membrane) fraction. **(B)** HeLa cells co-transfected with GCS-HA and GRASP55-GFP tagged constructs were fixed, premeabilized and stained with anti-HA antibody (Green) Scale bar, 10µm. **(C)** HeLa cells treated with indicated siRNAs and transfected with LCS-HA or co-transfected with GRASP55-GFP and LCS-HA were fixed, permeabilized and stained for LCS-HA (indicated in black). Spots of LCS-HA marked by black arrowheads (inset) indicate possible post-Golgi compartments. Scale bar, 10µm. **(D)** HeLa cells were fixed and processed for cryoimmunolabeling with an anti-GRASP55 antibody and anti-GOLPH3 antibody. Quantification of the distribution (LD normalised to the Golgi stack) of GRASP55 and GOLPH3 in peri-Golgi vesicles (carriers) is shown in the graph data are means ± SEM.

### Figure S11

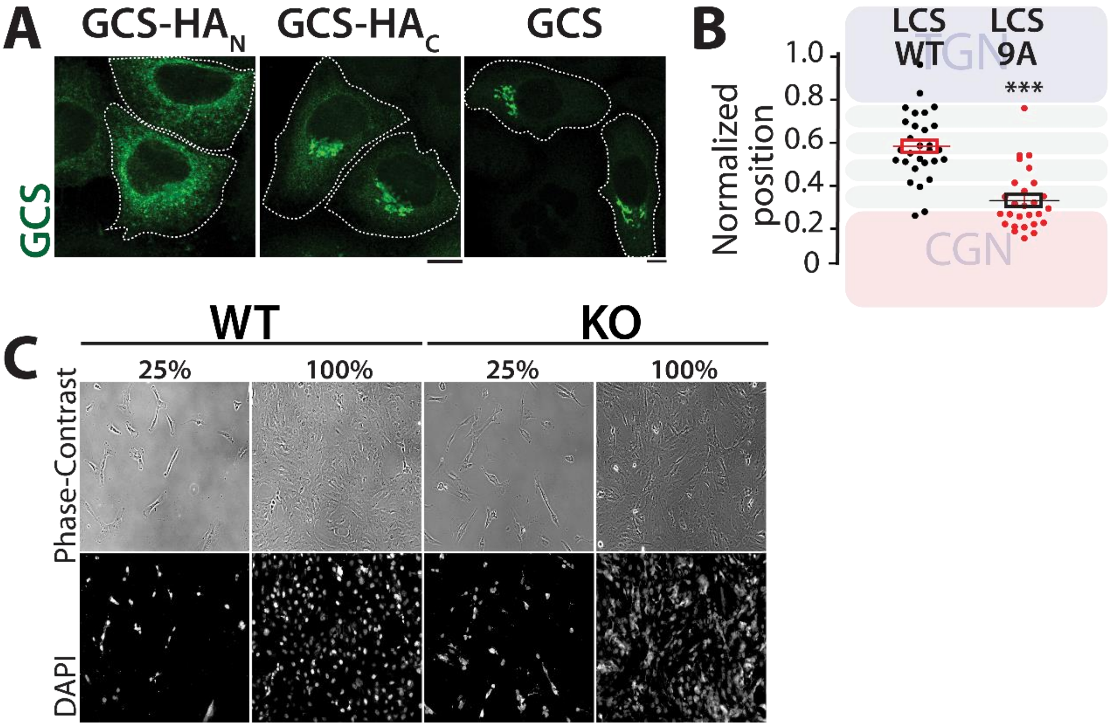


**Figure S11. Localization of GCS WT and mutants, sub-Golgi localization of LCS WT and mutant:**

**(A)** HeLa cells transfected with indicated GCS constructs were fixed, premeabilized and stained with anti-HA antibody (Green) Scale bar, 10µm. **(B)** HeLa cells were transfected with either WT LCS, or LCS9A, treated with nocodazole (33 µM) for 3 hours and labelled for enzymes, GM130, and TGN46. Line scan analysis was performed as in Fig.3A-B and the relative position of enzymes was quantitated and plotted. The data are mean ± SD; n> 30 stacks. ***p <0.001(Student's t test). **(C)** Fibroblasts were grown to indicated confluency, stained with DAPI and imaged. Scale bar, 100µm.

**Reagents:**

All reagents and chemicals were molecular biology grade. Methanol (Cat # 9093) and chloroform (Cat # 9180) were purchased from JT Baker, USA. Silica-gel high performance-TLC (HPTLC) plates (Cat #1055830001) was purchased from Merck, Germany. Fatty acid free Bovine Serum Albumin (Cat #A8806), HA Peptide (Cat #I2149), Brefeldin A (Cat #B7651) were purchased from Sigma-Aldrich, Germany. Protein A Sepharose CL-4B (Cat #17-0780-01) was purchased from GE Healthcare Life Sciences, USA. Anti-HA magnetic beads (Cat #88836) was purchased from ThermoFisher Scientific, USA. Sphingosine, [3-$^{3}$H]-, D-erythro>97% (Cat #NET1072050UC) was purchased from PerkinElmer, USA. Lipofectamine 2000 (Cat #11668027), Lipofecatime LTX with PLUS (Cat #15338100) were purchased from ThermoFisher Scientific, USA. TransIT-LT1 (Cat #MIR 2305) was purchased from Mirus, USA. RPMI 1640 (Cat #21875), DMEM (Cat #41965), DMEM/F-12 (Cat #11320033) and FBS (Cat #10437036) were purchased from Gibco/ThermoFisher Scientific, USA. BODIPY™ FL C5-Ceramide (Cat #D3521) was purchased from ThermoFisher Scientific, USA.  Protein A gold 15nm and Protein A gold 10nm were acquired from Cell Microscopy Core, UMC Utrecht. Biotin (Cat #29129) was purchased from Pierce, USA. Bacterial strains *E. coli* (DH5α) (Cat #18265017) and *E. coli* (BL-21-DE3) (Cat #C600003) were purchased from Thermo Fisher Scientific, USA. All the siRNAs and qPCR primers indicated in (**Tables S3 and S6)** respectively were purchased from Sigma-Aldrich, Germany.

**Cell lines:**

HeLa-M (human cervical cancer cells, female origin) were a kind gift from Prof. Paul Lehner, University of Cambridge. Wild type and GRASP55 knockout Human Fibroblasts (WI-26) were a kind gift from Markus Plomann, Institute for Biochemistry, University of Cologne. TALEN LCS-KO HeLa cell line (human cervical cancer cells, female origin) were kind gift from Kentaro Hanada, National Institute of Infectious Diseases, Japan. HeLa-M and GRASP55 KO HeLa cell lines (see below) were cultured in RPMI-1640 supplemented with 10% FCS. TALEN LCS-KO cell lines were cultured in DMEM supplemented with 10% FCS. Wild type and GRASP55 knockout Human Fibroblasts (WI-26) were cultured in DMEM: Nutrient Mixture F-12 (DMEM/F-12) supplemented with 10% FCS. All media were supplemented with 100U/ml penicillin/streptomycin and 2mM L-glutamine. All cells were grown in controlled atmosphere (5% $CO_2$ and 95% air) at 37 °C. Mycoplasma contamination was not observed in cell cultures as observed by DAPI staining. Cell cultures between 3-15 passages were used for the experiments and the cells were cultured to 80% confluence for the experiments unless indicated otherwise.

**Generation of GRASP55 Knockout cell lines by CRISPR-CAS9:**

To generate HeLa-M cell lines in which GRASP55 expression was abolished, we performed genome editing using CRISPR/Cas9 system. We obtained a pool of three plasmids each encoding guide RNA (gRNA) sequence designed to target GRASP55 coding sequence and pSpCas9 ribonuclease (Cat # SC-401106; Santa Cruz Biotechnology, Inc.). The list of gRNA sequences are reported in **Table S1**. These plasmids also encoded EGFP allowing positive selection of transfected cells. Hela-M cells were transfected with pooled plasmid for 48 hours, EGFP positive cells were isolated by FACS and single cells were sorted into each well of 96 well plates. The single cells were maintained in optimal culture conditions for 10 days, by replenishing fresh media every 48 hours. After 10 days, colonies formed from single cells were trypsinized and moved to 48 well plates and expanded to 6 well plates. Clones were collected and protein lysates were subjected to SDS-PAGE analysis and western blotting analysis using GRASP55 specific antibody to assess the presence of GRASP55 protein.

**Generation of expression constructs:**

<u>GCS-HA</u> was generated by inserting 9aa HA- tag into the C-terminal cytoplasmic tail of GCS between the two indicated Gly residues DPTISWRTGRYRLRC**GG**TAEEILDV. The following oligonucleotide primers used:

**F:**5'GCTGGAGAACTGGTCGCTACAGATTACGCTGTGGGTACCCATACGATGTTC CAGATTACGCTGGTACAGCAGAGGAAATCCTAGATGTATGATAACTCG-3'

**R:**5'CGAGTTATCATACATCTAGGATTTCCTCTGCTGTACCAGCGTAATCTGGAAC ATCGTATGGGTACCCACAGCGTAATCTGTAGCGACCAGTTCTCCAGC-3'.

<u>GCS Δ3C</u> (del of three C-terminal amino acids) was obtained by PCR amplifying GCS with the following primers to introduce a stop codon after the isoleucine residue at position -4 (DPTISWRTGRYRLRCGGTAEEILDV):

**F:** 5'-CATCGCGGATCCATGGCGCTGCTGGACCTG-3' and

**R:** 5’-GATCCGCTCGAGTTATCAGATTTCCTCTGCTGTACCAGCGTAATC-3’.

The PCR-generated fragment for both constructs was digested with EcoRI and XhoI (New England Biolabs, USA) and cloned into pcDNA4b-3xHA-expression vector. LCS9A Mutant (mutation of 9 amino acids to alanine in cytosolic tail of LCS) was generated by PCR amplification of LCS-WT-RUSH with following primers to introduce alanine into 9 amino acids (MAAAAAAAAAPRRSLLA):

**F:** 5’-CACAACCCGGGAGGCGCGCCATGGCAGCAGCAGCAGCAGCAGCAGCAGCA-3’

**R:** 5'- GCGAGCAGCGAGCGGCGCGGTGCTGCTGCTGCTGCTGCTGCTGCTGC -3'. The PCR-generated megaprimer was then annealed into LCS-WT-RUSH at 66ºC and extension at 72ºC for 25 cycles.

**Plasmids and siRNA transfection:**

HeLa-M cells, Wi-26 cells, LCS-KO cells and GRASP55 KO cells were transfected with plasmid vectors using TransIT-LT1 or Lipofectamine LTX reagents. A list of plasmids used in this study can be found in **Table S2**. Knockdown experiments for Hela-M cells were carried out using a pool of 4 siRNAs or 2 siRNAs using Lipofectamine 2000 according to manufacturer’s instructions (used at concentration of 100nM for the target gene). Expression or knockdown efficiencies (>85%) were checked after every experiment either by indirect immunofluorescence or immunoblotting or by qPCR analysis. A list of siRNA sequences used in this study can be found in **Table S3**.

**Cell lysis, western blotting and analysis:**

Cells were washed three times with ice cold PBS and lysed immediately at 4°C in RIPA lysis buffer (0.1% Triton X-100, 20mM Tris-HCl, pH 8.0, 0.1% SDS, 0.05% sodium deoxycholate, 150mM NaCl, 10mM $Na_3VO_4$, 40mM β-glycerophosphate, 10mM NaF) and complete protease/phosphates inhibitors (Roche). Cell lysates were clarified at 14,000 rpm for 10 min at 4°C to eliminate detergent insoluble pellet. To visualize GCS on SDS-PAGE, the protein lysates prepared as described above were treated with 250mM of DTT in sample buffer (62.5 mM Tris-HCL, pH 6.8, 2% SDS, 10% glycerol, 0.001% bromophenol, 125 mM dithiothreitol) at 37°C for 30 min. The lysate was immediately processed for SDS–PAGE and immunoblotting with antibodies. A complete list of primary and secondary antibodies are given in **Table S4**.

The western blots were then exposed to x-ray films and exposure time was varied to obtain optimal signal.

**Immunoprecipitation and peptide pull down assay**:

Total lysates were prepared using IP lysis buffer (150 mM NaCl, 25mM Tris-HCl pH 7.5, 1% Triton-X, 10mM $Na_3VO_4$, 40mM β-glycerophosphate, 10mM NaF and protease cocktail inhibitor from Roche). The protein concentrations were quantified using BCA Protein Assay kit (Pierce). 1mg of protein was used for precipitation with antibodies conjugated to either Protein A sepharose or magnetic dynabeads or to monomeric avidin beads (Pierce) in case of cytosolic peptides, and incubated at 4°C overnight. The beads were then washed 5 times in IP lysis buffer and bound proteins were subjected to SDS-PAGE and immunoblotted. The list of cytosolic peptides used in this study are reported in **Table S5**.

**Reconstitution of COPI vesicle formation:**

A two-stage incubation system was performed essentially as previously described (*1*). In brief, the first stage involved incubating prewashed (3M KCl) Golgi membrane (0.4 mg/ml) with coatomer (6 μg/ml), ARF1 (6 μg/ml), and 2 mM GTP in 500 μl of assay buffer (25 mM Hepes-KOH, pH 7.2, 50 mM KCl, 2.5 mM Mg(OAc)2, 1 mg/ml soybean trypsin inhibitor, 200 mM sucrose) for 15 minutes at 37°C. Afterwards, the Golgi membrane was pelleted (12,000 x g at 4°C for 10 minutes), and then resuspended in 100 μl of assay buffer for the second-stage incubation, which had ARFGAP1 (6 μg/ml) and BARS (6 μg/ml) added for 10 minutes at 37°C. Samples were then centrifuged for 10 minutes at 12,000 g and 4°C, with the pellet fraction containing the Golgi membrane and the supernatant fraction containing COPI vesicles.

**Protein Purification:**

Recombinant proteins were induced to express with 0.3 mM isopropyl β-D-thiogalactoside in BL21 (DE3) competent bacterial cells for 16h at 22 °C. Cells were harvested at 7000 rpm for 30 min at 4°C, lysed in lysis buffer (cold PBS with 1mM DTT, 1% triton X-100, phosphatase and protease inhibitors) and purified with Glutathione Sepharose 4B beads (GE Healthcare, USA). Recombinant GST-tagged proteins were eluted in elution buffer (50 mM Tris, 100 mM NaCl, 50 mM reduced gluthatione, pH 8.0); purified recombinant proteins have been concentrated and the buffer has been exchanged by using Vivaspin TURBO 4 filters (Sartorius, UK) at 4°C and 4,000 x g. The recombinant purified proteins were quantified by Bradford assay and were assessed for contaminants by SDS-PAGE.

**Isothermal titration calorimetry (ITC):**

ITC experiments were performed in a buffer containing 300 mM NaCl, 10 mM Bicine pH 8.5 and 1 mM DTT. Biotinylated Peptides were synthesized and delivered as lyophilized powder with a biotin moiety located at the N or C terminus (Charite Universitaetsmedizin Berlin, Germany). The peptides were dissolved in buffer, centrifuged at 14000 x g for 10 minutes and only the supernatant was used. The dissolved peptide concentrations were calculated based upon their absorbance at 280 nm and their corresponding molar extinction coefficient. Experiments consisted of titrations of 20 injections of 2 µL of titrant (peptides) into the cell containing GRASP domain protein at a 25-fold lower concentration. Typical concentrations for the titrant were around 2.5 mM for experiments depending on the affinity. Experiments were performed at 25 °C and a stirring speed of 1000 rpm on a MicroCal PEAQ-ITC (Malvern Panalytic). All data were processed using MicroCal PEAQ-ITC Analysis Software and fit to a one-site binding model after background buffer subtraction.

**GRASP55-GCS Peptide modelling:**

Model of GCS peptide was first built using backbone conformation of Golgin45 peptide as a template and fit into the same cleft of GRASP55, where Golgin45 peptide is located using COOT (*2*). This initial complex structure was further refined by the Docking2 module of Rosetta protocols through local docking search (*3-5*). The lowest energy model generated by this protocol was analyzed to probe the protein:peptide interaction. PRODIGY software was used to calculate the protein-protein binding energy (*6, 7*).

**Immunofluorescence and confocal microscopy:**

Indirect Immunofluorescence was performed as follows: cells were grown on 24mm coverslips, washed with PBS and fixed in 4% paraformaldehyde (Electron Microscopy sciences, Hatfield, USA) for 30 min at room temperature (RT). The cells were then permeabilized and blocked in blocking buffer (0.05% saponin and 0.5% BSA in PBS) for 30 min at RT followed by incubation with specific primary antibodies (see **Table S4**) for 1 hour at RT and washed with PBS. Cells were subsequently labelled with appropriate Alexa Fluor-conjugated secondary antibodies (**Table S4**). The coverslips were mounted using mowiol and images are acquired using confocal microscope Zeiss LSM700.

**Line Scan Analysis:**

The images were acquired with a pinhole set to 1 airy unit and under non-saturation conditions using a 63× objective (1.4 NA). Images were 8 bit with dimensions of 512x512 pixels and each pixel corresponded to an area of 132x132 $nm^2$. The line scan analysis was performed as described previously (*8*) using the Zen software system (Carl Zeiss). In brief, images of stacks stained for GM130, TGN46, and enzyme of interest tagged with HA were acquired as described earlier. Only cells with a moderate level of expression were considered for the analysis. Golgi stacks with clearly separated GM130-stained and TGN46-stained zones were identified and used for the analysis. A line was drawn in the middle of the stacks along the cis-trans direction, and the fluorescence intensity of each stained marker along this line was plotted. At least 30 stacks were examined per treatment, and a representative data is shown for analysis. The normalization of the distances was carried out by considering the start of the GM130 peak as 0, and the end of the TGN peak as 1. The images of Golgi stacks were processed using the "Image with Zoom" function of Metamorph 7.7.3.0 (Universal Imaging), for presentation.

**Cell profiler analysis:**

Control and GRASP55 KO cells were stained ShTxB-Cy3 and ChTxB-AlexaFluor 488 and images were acquired as described above using a 20X objective (NA 0.5). Images were 8 bit with dimensions of 512x512 pixels and each pixel corresponded to an area of 623x623 $nm^2$. For quantification experiments, 10-15 random fields were imaged with the same microscope settings (i.e. laser power and detector gain). The integrated intensity of fluorescence of each cell was calculated for each channel after the cells were segmented by Cell Profiler (*9*). Since only a fraction of the cells were positive for staining, the normalized intensity analysis was performed by taking top 10 percent of total cells in all the conditions and normalized on the background intensity.

**Flow cytometry analysis***:*

Control and GRASP55 KO cells were subjected to trypsin digestion, fixed with 4% paraformaldehyde, washed, and resuspended with PBS. Cells were incubated with bacterial toxins for 1 hr at 4°C. Then, cells were extensively washed with PBS and incubated with fluorescence-labeled secondary antibodies when required, or directly analyzed by BD FACS ARIAIII cell sorter (BD Biosciences). Cells incubated with secondary antibody alone, or unlabeled cells, were used as a negative control. The cell-surface expression of GSLs of selected cells were further analyzed in the gated region by BD FACS ARIAIII cell sorter (BD Biosciences). Bacterial toxins used are described in (Table S4).

**Electron microscopy:**

*Ultrastructural analysis of Golgi Morphology:* Briefly, control and GRASP55 knockout cells were grown in 35mm plastic dishes to 80% confluence. Cells were then fixed with 1% Glutaraldehyde (Electron Microscopy Sciences) in 0.2 M HEPES pH 7.3 at RT for 1h. The fixative was replaced with 1% BSA in PBS and cells were carefully detached using a plastic cell scraper, collected into microfuge tubes and centrifuged to obtain the pellet. All samples were then washed three times in 0.2 M HEPES pH 7.3 and post-fixed 30 minutes in 1% Osmium Tetroxide in the dark at 4°C in the same buffer. They were then washed three times in distilled water and post-fixed 25 minutes in 1% Osmium Tetroxide and 1.5% Potassium Ferrocyanide in the dark at room temperature in HEPES 0.2M pH 7.3. After washing three times in distilled water, they were stained with 0.5% uranyl acetate over night at 4°C. The pellets were dehydrated in graded steps of ethanol (50%, 70%, 90%, 100%), 2 times with 100% of acetone and embedded into Epon. Thin sections (60 nm) were cut on a Leica UC7 ultramicrotome and examined with 120 kV Philips Tecnai 12 Biotwin electron microscope (FEI, Eindhoven, The Netherlands) using a VELETA digital camera.

*Cryo-immuno EM:* Control or GRASP55 Knockout cells were transfected using indicated HA tagged construct for 16 hours. The cells were then fixed with 2%

formaldehyde and 0.2% glutaraldehyde in PHEM buffer (0.1M) pelleted by centrifugation, embedded in 12% gelatin, cooled on ice, and cut into 1-mm$^3$ cubes at 4 °C. The cubes were immersed in 2.3 M sucrose at 4 °C overnight, and then frozen in liquid nitrogen. Fifty-nanometre sections were cut with a diamond knife on a UC7 Leica cryo-ultramicrotome. The sections were picked up in a mix of 2% methyl cellulose and 2.3 M sucrose (1:1), as previously described (*10*), and collected on grids covered with Formvar-carbon supporting film (Electron Microscopy Sciences, PA, USA). The grids were first incubated with the rabbit anti-GM130 and/or mouse anti-HA polyclonal antibodies and then incubated with different sizes of Protein A gold (10 nm and 15 nm) to reveal antigen staining. After labelling, the sections were treated with 1% glutaraldehyde and embedded in methyl cellulose uranyl acetate for 10 min on ice. The excess of methyl cellulose uranyl acetate was removed and the sections were dried at room temperature before their analysis at 120 kV in a Philips Tecnai 12 Biotwin electron microscope (FEI, Eindhoven, The Netherlands) using a VELETA digital camera. The polarity of the Golgi stacks were by defined by compositional (cis Golgi marker GM130) parameter. For quantitation (performed with iTEM image acquisition software) cis, medial or trans Golgi were defined as previously described (*8*). Stacks with 4 cisternae were selected for the quantitation, Cis indicated the cis-most cisterna in the case of a stack with four cisternae. Trans was the last cisterna. Medial were two central cisternae. TGN was defined as the area in front of the trans cisterna upto a distance that equals the thickness of the Golgi stack. Vesicles were round profiles of 50–80 nm in diameter, present within 200 nm from the rims of the stack. The distribution of enzyme in the Golgi stack was expressed as the fraction of gold particles in each cisterna of the Golgi stack and TGN.

**Lipid Analysis**

HPLC-Mass Spectrometry:

Sphingolipids of control and GRASP55 KO cells were analysed by liquid chromatography tandem mass spectrometry (LC-MS/MS) as described earlier (*11*). Briefly, cells were washed twice with ice-cold PBS, and lipids were extracted with 2 ml ethyl acetate/isopropyl alcohol/water (60:30:10%, v/v/v) solvent mixture (*11*). The lipid extracts were analyzed with a Quantum Ultra triple quadrupole mass spectrometer connected to an Accela HPLC and Accela autosampler using a solvent gradient at the Lipidomics Core at Stony Brook University. Ceramides were identified through MRM analysis with soft fragmentation. Calibration curves were generated for each lipid and used for quantitative analysis of lipids in the samples. Inorganic phosphate (Pi) released from total phospholipids was measured by using a colorimetric method. Briefly, following a lipid extraction by using the Bligh and Dyer method, samples were dried under a stream of nitrogen gas. Then, a mixture of sulfuric and hydrochloric acid were added to the sample to ash the organic content. Samples were heated overnight at 160ºC. Then, water, ammonium molybdate and ascorbic acid were subsequently added to samples and incubated for 30 min at 45ºC. The absorbance was measured at 600 nm by using a Spectramax M5 plate reader. A calibration curve was created and utilized to quantify the inorganic lipid content for each sample and used as normalization for relative quantification of sphingolipids.

Radioactive pulse-chase assay to monitor SL biosynthesis:

Radioactive pulse chase assay was performed as described earlier (*12*). Briefly, HeLa-M cells or human fibroblasts wi-26 or GRASP55 KO cells were pulse labelled for 2 hours with [$^3$H]-sphingosine (final concentration of 0.1 µCi/mL; Perkin Elmer) in serum-free DMEM containing 1 % fatty acid free BSA followed by a chase for indicated times in complete media. Lipids were then extracted from the cells, resuspended in chloroform, spotted onto silica-gel high performance-TLC (HPTLC) plates (Merck, Germany), resolved using a mixture of chloroform, methanol and water (65:25:4 v/v/v) and quantified using GINA® (Raytest, Germany) software analysis.

MALDI-MS

Total lipid extracts were prepared using a standard MTBE protocol followed by a methylamine treatment for sphingo- and glycosphingolipids analysis by mass spectrometry. Briefly, cell pellet was resuspended in 100 µL $H_2O$. 360 µL methanol and 1.2 mL of MTBE were added and samples were placed for 10 min on a vortex at 4 °C followed by incubation for 1 h at room temperature on a shaker. Phase separation was induced by addition of 200 µL of $H_2O$. After 10 min at room temperature, samples were centrifuged at 1000 g for 10 min. The upper (organic) phase was transferred into a glass tube and the lower phase was re-extracted with 400 µL artificial upper phase [MTBE/methanol/H2O (10:3:1.5, v/v/v)]. The combined organic phases were dried in a vacuum concentrator. Lipids where then resuspended in 500 µL of $CHCl_3$ and divided in two aliquots for a further methylamine treatment. 500 µL of freshly prepared monomethylamine reagent [methylamine/H2O/n-butanol/methanol (5:3:1:4, (v/v/v/v)] was added to the dried lipid extract and then incubated at 53 °C for 1 h in a water bath. Lipids were cooled to room temperature and then dried. The dried lipid extract was then extracted by n-butanol extraction using 300 µL water-saturated n-butanol and 150 µL $H_2O$. The organic phase was collected, and the aqueous phase was re-extracted twice with 300 µL water-saturated n- butanol. The organic phases were pooled and dried in a vacuum concentrator. Lipids were then resuspended in 500 µL of $CHCl_3$ and analyzed by MALDI-MS. 30 mg/mL 2,5-DHB was freshly prepared in acetonitrile/water solution (50:50 v/v) with 0.1% TFA. An equivalent volume of sample solution (50 µL) was then mixed with matrix before deposition on the MALDI target. All mass spectrometry analysis for the identification of lipids (m/z 600-1800) were obtained using an Applied Biosystems 4800 MALDI-TOF/TOF mass spectrometer equipped with a 200 Hz tripledfrequency Nd:YAG pulsed laser with 355 nm wavelength. Measurements were performed in positive ion reflection mode at an accelerating potential of 20 kV. Each mass spectra were obtained by applying a laser energy of 4600 watts/cm$^2$, averaging 4000 single laser shots/spectrum.

**FRAP assay:**

Control and GRASP55 (#2) knockout cells were plated on 35-mm glass bottom microwell dishes (MatTech, USA) and transfected with CERT-YFP (1 μg) for 16 hours. FRAP was performed using Zeiss LSM 710 confocal microscope equipped with an environmental control system set to 37°C and 5% $CO_2$. FRAP experiments were performed by bleaching CERT-YFP in the Golgi area (50 bleaching iterations) followed imaging every 7.5 seconds for 4 minutes (2% laser power). Recovery of fluorescence was calculated as ratio of Golgi intensity and that of total cell after background correction.

**Ceramide Transport assay:**

Control and GRASP55 knockout cells were grown on coverslips to 80% confluence. Cells were washed with DMEM-HEPES containing 10% FCS for 3 times and incubated with BODIPY ceramide (5μM) in fatty acid free BSA for 30 minutes on ice. After incubation, cells were washed with DMEM-HEPES for 3 times and followed by imaging. The laser conditions used were - 488nm (excitation) and 620nm (emission) and fluorescence was recorded every 2 minutes for a period of 20 minutes. The fluorescence of bodipy-Ceramide in the Golgi was determined followed by normalization to the maximum.

**RNA extraction and Real Time PCR:**

Total RNA was extracted from HeLa-M cells (Control and GRASP55 Knockout) using the RNeasy Mini Kit (Qiagen) according to manufacturer's instructions. The yield and the integrity of RNA were determined by spectrophotometer NanoDrop 2000c (Thermo Scientific) and by TAE agarose gel electrophoresis, respectively. RNA (1 μg) was reverse transcribed using QuantiTect Reverse Transcription Kit (Qiagen) according to the manufacturer's instructions and subjected to qPCR with gene specific primers (**Table S6)** in the presence of LightCycler®480 SYBR Green I Master Mix (Roche, Switzerland) on a LightCycler®480 II detection system (Roche, Switzerland). Analyses were carried out on biological triplicate samples for each experiment and they were processed separately. The thermal profile consisted of 10 min at 95°C pre-incubation, and 40 cycles at 10sec at 95°C, 10sec at 60°C and 10sec at 72°C. The qPCR data were normalized to the average of the reference gene human hypoxanthine-guanine phosphoribosyltransferase 1 (HPRT1). The fold changes in the relative quantifications were calculated according the ΔΔCt method. All the commercial kits used are reported in (**Table S7**).

**Experimental conditions for RUSH experiments:**

LCS-KO cells were transiently transfected with LCS or LCS9A constructs at 37°C in absence of biotin for 16 hours, to localize the proteins to endoplasmic reticulum (ER). Their ER exit was promoted by addition of biotin (40 µM) for 6 hours. For pulse chase assay experiments (**Fig.4C**) biotin was replaced every 3 hours and maintained throughout chase period for a total of 24 hours.

**Edu proliferation Assay:**

WT and GRASP55 KO fibroblasts were grown to 100% confluence. EdU (10 µM) was added to the culture medium and cells were incubated overnight. After Edu incubation, cells were washed with 3 mL of 1% BSA in PBS. Cells were fixed by addition of 100 µL of Click-iT® fixative for 15 minutes. After fixation, cells were washed with 3 mL of 1% BSA in PBS, centrifuged and supernatant was discarded. Pellet was dislodged by slight agitation and were premeabilized by addition of 100 µL of 1X Click-iT® saponin-based permeabilization buffer. After the permeabilization, Click-iT® reaction cocktail containing PBS, CuSO4, Fluorescent dye azide ( Alexa Fluor® 488 azide) was added to cells and incubated for 30 mins. After labelling cells were washed with PBS and were analysed by BD FACS ARIAIII cell sorter (BD Biosciences).

**Foci formation assay:**

Wild type and GRASP55 KO wi-26 fibroblasts were seeded onto 10cm petri dishes (Corning, USA) at the confluence of $2x10^6$ and grown for 10-12 days. Growth media was replaced every 48 hours and for treatment with glycosphingolipid inhibitor, media was replaced every 48 hours with1-phenyl-2-decanoylamino-3-morpholino-1-propanol (PDMP) (2 µM). Cells were fixed with methanol on ice for 10 mins and foci were stained using crystal violet (0.1 % w/v) for 10 min, washed extensively with water and counted using ImageJ software.

**Statistics:**

Error bars correspond to either standard deviation (SD) or standard error of mean (SEM) as indicated in figure legends. All the statistical evaluations were done using GraphPad Prism built-in tests (unpaired two-tailed Student's t-tests) and as indicated significance values are all marked as follows * $P< 0.05$, ** $P< 0.01$, and ***$P < 0.001$ (*ns*, not significant). All the measurements reported are from distinct samples.

**Data availability Statement:**

Source data are provided with the manuscript and if not, they are available from the corresponding author upon reasonable request.

#### Supplementary Tables

(**Table S1**) gRNAs used in this study to generate GRASP55 knockout cell lines.

| gRNA | Sequence | | |
|---|---|---|---|
| #1 GUIDE RNA | CAGTCACACCAAGTAACCTG | Santa Cruz Biotechnology, Inc | Cat # SC-401106; |
| #2 GUIDE RNA | TTGCTTCATGTGTTTCGATA | Santa Cruz Biotechnology, Inc | Cat # SC-401106 |
| #3 GUIDE RNA | CTATAGATAAGCATCTTTAC | Santa Cruz Biotechnology, Inc | Cat # SC-401106 |

(**Table S2**). List of recombinant DNA used in this study

| Recombinant DNA | Source | |
|---|---|---|
| GRASP55-EGFP | Kind gift from Yanzhuang Wang | *N/A* |
| GB3S | Kind gift from Antonella De Matteis | *N/A* |
| GM3S | Kind gift from Antonella De Matteis | *N/A* |
| LCS | Kind gift from Antonella De Matteis | *N/A* |
| GCS-HAC | Kind gift from Giovanni D’angelo | *N/A* |
| SMS1 | Kind gift from Giovanni D’angelo | *N/A* |
| CERT | Kind gift from Antonella De Matteis | *N/A* |
| PEGFP | Addgene | Cat # 6085-1 |
| Sar1-GTP | Kind gift from Rainer Pepperkok | *N/A* |
| ARF1-GTP | Kind gift from Antonella De Matteis | *N/A* |
| Str-KDEL_B4GALT5-SBP-EGFP (referred as LCS WT) | This study. | *N/A* |

| | | |
|---|---|---|
| Str-KDEL_B4GALT5-SBP-EGFP (referred as and LCS9A) | This study. | *N/A* |
| GCS-HA$_N$ | This Study (custom made from genescript) | *N/A* |
| GCS | This Study | *N/A* |
| GCS Δ3C | This Study | *N/A* |
| pGEX6p-1GRASP55 N | Kind gift from Yanzhuang Wang | *N/A* |
| pGEX4T-1GRASP55 C | Kind gift from Yanzhuang Wang | *N/A* |
| pGEX6p-1GRASP55 G97D | This Study | *N/A* |
| GRASP55-His tag | Kind gift from Antonino Colanzi | *N/A* |
| GRASP65-His tag | Kind gift from Antonino Colanzi | *N/A* |

(**Table S3**) siRNA sequences used in this study to downregulate indicated human gene expression

| **Human Gene** | **Accession Number** | **siRNA Sequence** |
|---|---|---|
| Giantin/GOLGB1 | NM_004487 | #1 5’-GAACUAGAGUCUCGGUAUA-3’<br>#2 5’-UAAGAGAAUUGCAACCUAA-3’<br>#3 5’-GUACACAGGUUAAGUGCUU-3’<br>#4 5’-GAAGGUCUGUGAUACUCUA-3’ |
| GM130/GOLGA2 | NM_004486 | #1 5’-GGACAAUGCUGCUACUCUA-3’<br>#2 5’-AGAAGGAGGUGCUGCAUAA-3’<br>#3 5’-GAAUAUCAGCAGAGGAAUA-3’ |

| | | |
|---|---|---|
| | | #4 5’-UUGUAAAGCUGACUAAUGA-3’ |
| GRASP 65/GORASP1 | NM_031899 | #1 5’-GAUCUCUACCACAGAAUAA-3’<br>#2 5’-CUGGAGGUGUUCAAUAUGA-3’<br>#3 5’-GAGGACUUCUUUACGCUCA-3’<br>#4 5’-GGACGUGUCGGGAAUUUCU-3’ |
| GRASP 55/GORASP2 | NM_015530 | #1 5’-GGAGUGAGCAUUCGUUUCU-3’<br>#2 5’-GUAAACCAGUCCCUCACUU-3’<br>#3 5’-GACCACACAGUGAUUAUAU-3’<br>#4 5’-UGUCGAGAAGUGAUUAUUA-3’ |
| GMAP210/TRIP11 | NM_004239 | #1 5’-GGACAUUACUAAAGAGUUA-3’<br>#2 5’-GGGCAAGACUGGAGAGUUA-3’<br>#3 5’-GAACUUAAGGAGCAUAUUA-3’<br>#4 5’-GGACUUUGGUGAUAUAAUU-3’ |
| CERT | NM_001130105 | #1 5’-GAAGAUGACUUUCCUACAAUU-3’<br>#2 5’-GAAGUUGGCUGAAAUGGAAUU-3’<br>#3 5’-GCGAGAGUAUCCUAAAUUUUU-3’<br>#4 5’-UCAAAGGGAUAAAGUGGUAUU-3’ |
| Golgin 97/GOLGA1 | NM_002077 | #1 5’- AAGAUCACAGCCCUGGAACAA -3’<br>#2 5’- AAGUGCUUCUCCAGAAAGAGC -3’ |
| Golgin 45/BLZF1 | NM_003666 | #1 5’-UGUGAUGUAUGGCGAAGUA-3’<br>#2 5’-GAAUAUUAGUUCCCAAAGC-3’<br>#3 5’-GAACGUCUAGCCCGUGAGA-3’ |

| | | |
|---|---|---|
| | | #4 5’-GAACAGUUAGAACGUAUGU-3’ |
| GCC1 | NM_024523 | #1 5’-GGACUUGGAGCUUAGGUUA-3’<br>#2 5’-GGAGAAGGCUACUGCACUC-3’ |
| GCC2 | NM_181453 | #1 5’-UCAAAGAGAUACCAUGUUA-3’<br>#2 5’-GAGCAGAGUUGAUACUAUU-3’ |
| Golgin 160/GOLGA3 | NM_005895 | #1 5’-GCAAACAGCCCGUGGGAAA-3’<br>#2 5’-GAGAUGAAGACCAAACAUA-3’ |
| Golgin 245/GOLGA4 | NM_001172713 | #1 5’-GAACUAACCUGUCAGAUUU-3’<br>#2 5’-GAGAACAGAUUCACAAUUU-3’ |
| GOLPH3 | NM_022130 | 1# 5’-AAAUGAUGUGUAACCCUCGCGGUCC-3’<br>2 #5’-AAUCCAGAUGAUAUACAGUCAUUCC -3’<br>6 #5’-GGAGAGGAAGGUUACAACUA-3’<br>7# 5’-UCAAGGACCGCGAGGGUUA -3’ |
| GOLIM4 | NM_014498 | #1 5’-GAACACAGAUCAAGAUUAG-3’<br>#2 5’-GCAGAUGACCCUAAUAAUC-3’ |
| GOPC | NM_020399 | #1 5’-GGCCUUGGCAUUUCAAUUA-3’<br>#2 5’-GGACAUCGUUACCGUUUGU-3’ |

| | | |
|---|---|---|
| GORAB | NM_152281 | #1 5’-GCAGAGACCAUGAAACUAA-3’<br>#2 5’-CAGCAAAGCUAGAUAUACA-3’ |
| FAPP2 | NM_001197026 | #1 5’-GAGAUAGACUGCAGCAUAUUU-3’<br>#2 5’-GAAUUGAUGUGGGAACUUUUU-3’<br>#3 5’-GAAAUCAACCUGUAAUACUUU-3’<br>#4 5’CCUAAGAAAUCCAACAGAAUU-3’ |
| AllStars Negative Control siRNA | Qiagen | Cat #SI03650318 |

(**Table S4**) List of antibodies used in this study for immunofluorescence (IF), Western Blotting (WB), cryo-Electron Microscopy (cryo-EM).

| Antibodies | SOURCE | IDENTIFIER |
|---|---|---|
| Monoclonal Mouse GMAP210 | BD Biosciences | Clone 15<br>Cat # 611712<br>RRID:AB_399190 |
| Monoclonal Mouse GM130 | BD Biosciences | Clone 35<br>Cat # 610822<br>RRID: AB_398141 |
| Polyclonal Rabbit GIANTIN | Abcam | Cat # ab24586<br>RRID: AB_448163 |
| Monoclonal Mouse GRASP65 | Santa Cruz Biotechnology | Clone D12<br>Cat # sc-374423<br>RRID: AB_10991322 |
| Polyclonal Rabbit GOLPH3 | Abcam | Clone |

| | | |
|---|---|---|
| | | Cat # ab98023<br><br>RRID: AB_10860828 |
| Polyclonal Rabbit B4GALT1 | Sigma | Cat # HPA010807<br><br>RRID: AB_1078254 |
| Monoclonal Mouse GAPDH | Santa Cruz Biotechnology | Clone 6C5<br><br>Cat #sc-32233<br><br>RRID:AB_627679 |
| Polyclonal Rabbit GRASP55 | Novus Biologicals | Cat #NBP1-89747<br><br>RRID:AB_11024556 |
| Monoclonal Mouse HA-Tag | Biolegend/Covance | Clone 16B12<br><br>Cat #MMS-101P<br><br>RRID:AB_10063630 |
| Monoclonal Rabbit HA-Tag | Cell Signaling Technology | Clone C29F4<br><br>Cat #3724 |
| Polyclonal Sheep anti-human TGN46 | BioRad/AbD-Serotec | Cat #AHP500G<br><br>RRID:AB_323104 |
| Monoclonal Mouse anti-β ACTIN | Sigma | Clone AC-74<br><br>Cat #A2228<br><br>RRID:AB_476697 |
| Monoclonal Mouse GFP | Abcam | Cat #ab6556<br><br>RRID:AB_305564 |
| GST | This study | N/A |
| Anti-polyHistidine | Sigma | Clone HIS-1<br><br>Cat #H1029, |

| | | |
|---|---|---|
| | | RRID:AB_260015 |
| ShTxB-CY3 | Dr. Ludger Johannes | *N/A* |
| ChTxB-AlexaFluor 488 | Invitrogen | Cat # C-22841 |
| Anti-mouse, donkey Alexa Fluor 488 | ThermoFisher Scientific | Cat #A-21202 RRID:AB_141607 |
| Anti-mouse, donkey Alexa Fluor 568 | ThermoFisher Scientific | Cat #A10037 RRID:AB_2534013 |
| Anti-mouse, goat Alexa Fluor 633 | ThermoFisher Scientific | Cat #A-21052 RRID:AB_141459 |
| Anti-rabbit, donkey Alexa Fluor 488 | ThermoFisher Scientific | Cat #A-21206 RRID:AB_141708 |
| Anti-rabbit, donkey Alexa Fluor 568 | ThermoFisher Scientific | Cat #A10042 RRID:AB_2534017 |
| Anti-rabbit, goat Alexa Fluor 633 | ThermoFisher Scientific | Cat #A-21070 RRID:AB_2535731 |
| Anti-sheep, donkey Alexa Fluor 633 | ThermoFisher Scientific | Cat #A-21100 RRID:AB_2535754 |
| Anti-goat, donkey Alexa Fluor 568 | ThermoFisher Scientific | Cat #A-11057 RRID:AB_142581 |

(**Table S5**) List of cytosolic peptides used in this study for immunoprecipitation assays.

| **Cytosolic Tail Peptides** | | |
|---|---|---|
| B4GALT-1 cytosolic tail peptide<br>Biotin- MRLREPLLSGSAAMPGA -COOH | This study | N/A |
| LCS cytosolic tail peptide<br>Biotin- MRARRGLLRLPRRSLLA -COOH | This study | N/A |
| LCS Mut_1 cytosolic tail peptide<br>Biotin- MRARRGLLAAAAASLLA -COOH | This study | N/A |
| LCS Mut_2 cytosolic tail peptide<br>Biotin-AAAAAGLLRLPRRSLLA -COOH | This study | N/A |
| LCS Mut_3 cytosolic tail peptide<br>Biotin-MRARRAAAAAPRRSLLA -COOH | This study | N/A |
| GCS Cytosolic tail peptide<br>Biotin-DPTISWRTGRYRLRCGGTAEEILDV-COOH | This study | N/A |
| GCS Δ 3C cytosolic tail peptide<br>Biotin- PTISWRTGRYRLRCGGTAEEI -COOH | This study | N/A |

(**Table S6**) Primers used in this study to determine mRNA levels of indicated gene.

| Human | Forward Primer | Reverse Primer |
|---|---|---|
| GCS | 5’-TTCGGGTTCGTCCTCTTC-3’ | 5’-GCTTGCTATAAGGCTGTTTGTC-3’ |
| LCS | 5’-CAATCGGTGCTCAGGTTTATG-3’ | 5’-GGTTTCACTGTGGTTCAAGTC-3’ |
| GB3S | 5’-GATCCCCACCTCTCTGCAAT-3’ | 5’-TTGGACATGGTATCCCCAGA-3’ |
| GM3S | 5’-TGGTTATTGGAAGCGGAGG-3’ | 5’-TCTGAATATCCCTCAACTGGT-3’ |
| FAPP2 | 5’-ACATCAGGATCCGATTGAGA-3’ | 5’-ATGCACCTTCTGGATGTGTT-3’ |
| CERT | 5’-TGTGGATCATGACAGTGCTC-3’ | 5’-ATTTCCTGGTTTCCCTCTGG-3’ |
| SMS1 | 5’-GCATTTCAACTGTTCTCCGAAG-3’ | 5’-GATAGACAAGCCACCTCCAG-3’ |

| | | |
|---|---|---|
| HPRT1 | 5'-AGCTTGCTGGTGAAAAGGAC-3' | 5'-GTCAAGGGCATATCCAACAAC-3' |
| GM2S | 5'-CAGAAACAAGTCCGAGCTATTGA-3' | 5'-GAGGGGCTGAACTTCCACAC-3' |
| GM1S | 5'-CAACCTCACCCTAAAGACCC3' | 5'-AGGGACGTTGACATACACATC-3' |

(**Table S7**) All the commercial kits used for cDNA extraction and mRNA analysis.

| **Commercial assays and kits** | | |
|---|---|---|
| RNA easy mini | Qiagen | Cat #74106 |
| QuantiTect Reverse Transcription Kit<br>Print | Qiagen | Cat #205311 |
| SYBR™ Green PCR Master Mix | ThermoFisher Scientific | Cat #4309155 |
| Click-iT™ EdU Alexa Fluor™ 488 Flow Cytometry Assay Kit | ThermoFisher Scientific | Cat #C10420 |

(**Table S8**) All the Software used for analysis and figure preparation.

| **Softwares** | | |
|---|---|---|
| ImageJ | NIH | https://imagej.nih.gov/ij/ |
| MetaMorph | Molecular Devices | https://www.moleculardevices.com/systems/metamorph-research-imaging |
| Prism | Graphpad | https://www.graphpad.com/scientific-software/prism/ |
| Zen Lite | Carl Zeiss | https://www.zeiss.com/microscopy/int/products/microscope-software/zen-lite.html |
| Adobe Illustrator | Adobe | www.adobe.com/products/illustrator/free-trial-download.htm |
| Adobe Photoshop | Adobe | www.adobe.com/products/photoshop.html |
| Soft Imaging service (Electron microscope) | Olympus | www.olympus-sis.com/corp/2256.htm |
| iTEM | EMSIS GmbH | https://www.emsis.eu/products/software/item/ |
| Cell profiler | Cell Profiler Analyst | https://cellprofiler.org/releases/ |
